# Team flow is a unique brain state associated with enhanced information integration and neural synchrony

**DOI:** 10.1101/2020.06.17.157990

**Authors:** Mohammad Shehata, Miao Cheng, Angus Leung, Naotsugu Tsuchiya, Daw-An Wu, Chia-huei Tseng, Shigeki Nakauchi, Shinsuke Shimojo

## Abstract

Team flow occurs when a group of people reaches high task engagement while sharing a common goal as in sports teams and music bands. While team flow is a superior enjoyable experience to individuals experiencing flow or regular socialization, the neural basis for such superiority is still unclear. Here, we addressed this question utilizing a music rhythm task and electroencephalogram hyper-scanning. Experimental manipulations held the motor task constant while disrupted the hedonic musical correspondence to blocking flow or occluded the partner’s body and task feedback to block social interaction. The manipulations’ effectiveness was confirmed using psychometric ratings and an objective measure for the depth of flow experience through the inhibition of the auditory-evoked potential to a task-irrelevant stimulus. Spectral power analysis revealed higher beta/gamma power specific to team flow at the left temporal cortex. Causal interaction analysis revealed that the left temporal cortex receives information from areas encoding individual flow or socialization. The left temporal cortex was also significantly involved in integrated information at both the intra- and inter-brains levels. Moreover, team flow resulted in enhanced global inter-brain integrated information and neural synchrony. Thus, our report presents neural evidence that team flow results in a distinct brain state and suggests a neurocognitive mechanism by which the brain creates this unique experience.

**Data Availability:** All data and analysis codes used in the preparation of this article are available at https://osf.io/3b4hp.

## INTRODUCTION

Flow state, or getting into the zone, is a psychological phenomenon that develops when there is a balance between the skills of the individual and the challenge of the task, clear goals, and immediate feedback (1, 2). Flow is accompanied by intense task-related attention, effortless automatic action, a strong sense of control, and a reduced sense of external and internal awareness and sense of time (2). Flow is intrinsically rewarding and have a positive effect on several life experiences (1–4). Although flow can develop during an individual activity, it is more common to develop during a group activity. There is a growing interest in studying flow in group activities, group flow, within several fields including psychology, sociology, organizational behavior, and business fields; hence, it has been studied in a plethora of contexts concerning sports, music, education, work, and gaming (5–10). Group flow, such as classroom students, audience in a concert or a group of motorcycle drivers, is believed to be a collective phenomenon, not a simple aggregation of individual flow experiences, showing augmented positive effects, including enhanced creativity, productivity, or emotions (5–10). Team flow is a specific case of group flow where the group forms a team characterized by a common purpose, complementary skills, performance goals, strong commitment, and mutual accountability (11, 12). The positive subjective experience during team flow, as in sports teams, music ensembles, dance squads, business teams, or teams in video gameplay, is superior to ordinary socialization or individuals experiencing flow (5, 9, 13). Flow and socialization are two disparate subjective experiences; in other words, acting in a social context is not necessarily sufficient to get into the flow state, and vice versa.

The neural mechanisms of both individual flow and socialization experiences have been studied in isolation. For social information processing, several networks have been implicated: social perception, empathy, mentalization, and action observation networks have been identified as partially overlapping brain regions with the central role played by the amygdala, the anterior cingulate cortex (ACC), the prefrontal cortex (PFC), and the inferior frontal gyrus (IFG) and the inferior and superior parietal lobule (IPL/SPL), respectively (14–19). Meanwhile, several studies of individual flow have shown increased activity of the IFG and the IPL/SPL, and a decreased activity of the PFC (20–24). There are concordant and discordant overlaps between the brain regions involved in these two experiences. For example, both experiences activate IFG and IPL/SPL. On the other hand, the reported inhibition of the PFC activity during individual flow is discordant with the reported social perception and mentalization information processing. Given the nature of the complex overlap between the brain regions involved in individual flow and socialization, a unique interaction among these brain areas might emerge during the team flow experience. While, phenomenologically, the experience of team flow is subjectively more intense than individual flow and ordinary socialization, the underlying neurocognitive mechanism is still unclear. The current study aimed to directly examine the possible distinct neural activity patterns, emerging at both the intra-brain and inter-brains levels during team flow.

## RESULTS

### Behavioral establishment and subjective assessment of team flow

To address these questions, we established a behavioral paradigm where a pair of prosocial highly-skilled participants are engaged in playing a popular music rhythm game. The game’s task is to respond by tapping a touch screen at the moment animated visual cues reach a designated area with instantaneous positive feedback. The feedback and the well-designed cues give the impression of playing a musical instrument inducing the game’s positive experience and a state of flow. Each pair of participants, matched in skill level and song preference, played as a team through splitting the tapping area and sharing in completing the task with a common goal of getting the best score for the team (Fig. 1a and Movie S1). In the meanwhile, we simultaneously recorded their brain activities using electroencephalogram (EEG).

**Figure 1.**
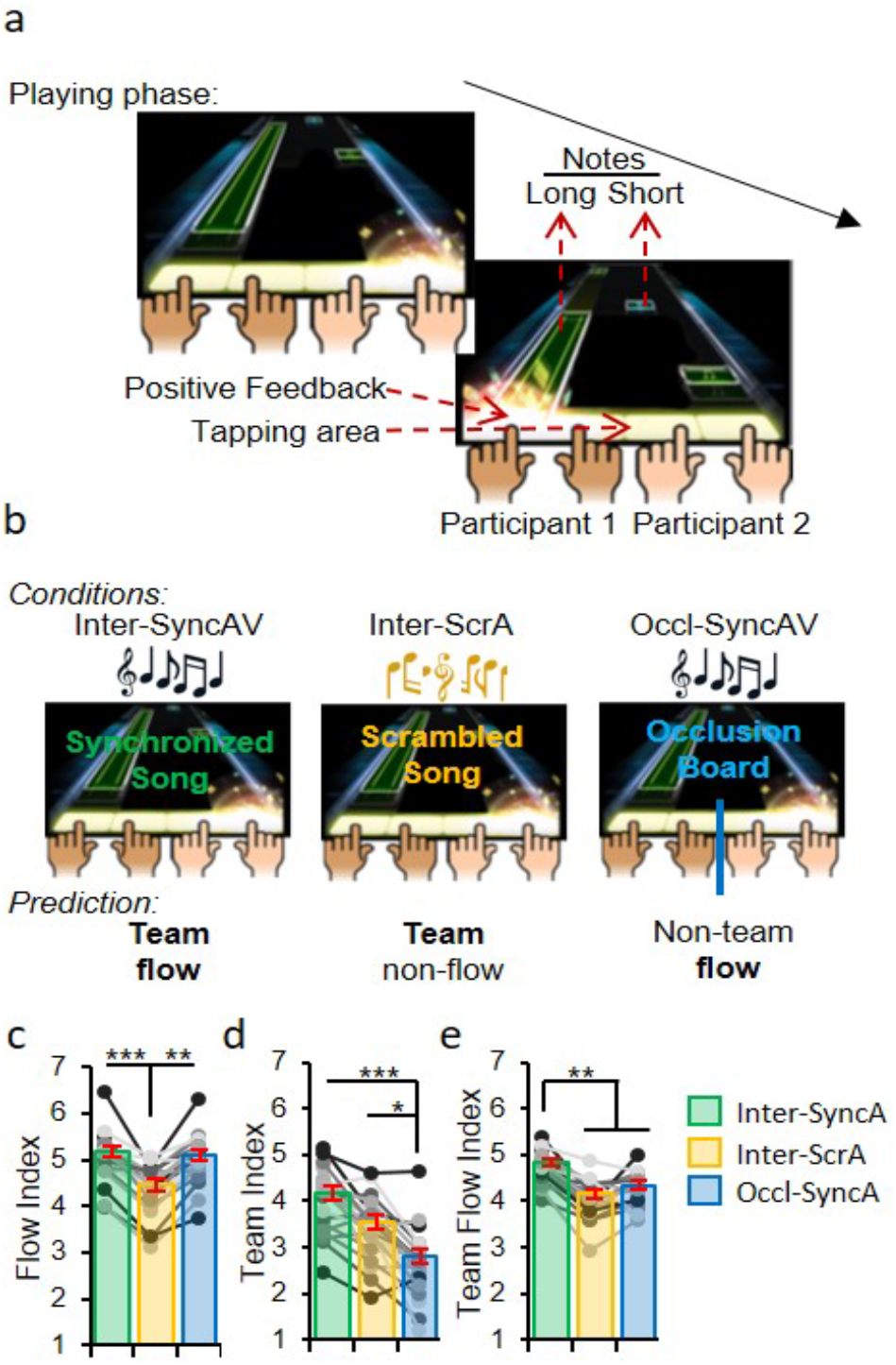
Behavioral establishment and subjective assessment of team flow. **a**, Diagram of the finger-tapping music rhythm task. Participants must tap when animated cues/notes moving from the top of the screen reach the tapping area. **b**, Manipulations (see Table S1): team flow is predicted when the participants are playing the original song with the auditory-visual input synchronized (Inter-SyncA). Flow is interrupted through scrambling the music (Inter-ScrA). Team interaction is interrupted through hiding the partner’s body and feedback using an occlusion board between participants (Occl-SyncA). **c-e**, Subjective rating indices as a measure of flow experience, interpersonal interaction or interpersonal flow (see Fig. S2). Friedman test with Dunn’s post-hoc. * P < 0.05, ** P < 0.01, *** P < 0.001. Error bars represent mean ± s.e.m.; n = 20.

In the primary experimental condition, teams were playing in an open interpersonal setting, and the visual tapping cues were accompanied by the synchronized audio music track (Interpersonal synchronized audio condition, or Inter-SyncA) (Fig. 1b and Table S1). This condition was designed to maximize the team flow experience. To disrupt the flow experience, we manipulated the intrinsic reward/enjoyment dimension of flow by scrambling the game’s music, hence, interrupting the sense of immersion and continuity of the game (Inter-ScrA) (Fig. 1b and Table S1). To interrupt the social (team) interactions, we used an occlusion board between the two participants that occluded the partner’s whole bodies and feedback while leaving all of the cues visible to both players (Occl-SyncA) (Fig. 1b, Fig. S1a, and Table S1). We expected that the three experimental conditions, namely, Inter-SyncA, Inter-ScrA, and Occl-SyncA, would provide optimal manipulations to produce team flow, team non-flow, or non-team flow experiences, respectively.

To validate our manipulations, participants performed psychometric ratings after each trial, which were indexed along the dimensions of flow and team interactions (Fig. S2). As expected, the subjective experience of flow, assessed by a flow index, significantly decreased in the Inter-ScrA condition less than the other two conditions (Friedman Test, chi-square = 25.0, p < 0.0001; Fig. 1c and Fig. S2). The subjective positive team interactions, assessed using a team index, significantly decreased in the Occl-SyncA condition less than the other two conditions (Friedman Test, chi-square = 25.291, p < 0.0001; Fig. 1d and Fig. S2). The subjective experience of team flow, assessed using the team flow index, was significantly higher in the Inter-SyncA condition more than the other two conditions (Friedman Test, chi-square = 26.8, p < 0.0001; Fig. 1e and Fig. S2). Also, there were no differences in the participants performance across conditions (one-way ANOVA, F(2,27) = 0.02437, p = 0.976), excluding possible neural differences caused by motor responses. Collectively, the results of the psychometric assessment confirmed effectiveness of our manipulations to achieve the aimed state, while maintaining minimal changes in other factors, such as task difficulty, sensory input, and motor response (Table S1).

### The depth the flow state was confirmed by neurophysiological measure

To provide a more objective evidence for the flow state, we established a novel neurophysiological measure of flow. We utilized the intense task-related attention, and the reduced sense of external awareness dimensions of flow (2), and the well-known effect of selective attention on the auditory-evoked potential (AEP) (25). During each trial, we presented task-irrelevant beeps to the participants. The strength of the resultant AEP to the task-irrelevant beeps should be inversely proportional to the immersion in the game and thus constitute an objective measure for flow (Fig. 2 and Fig. S3). As expected, the mean AEP response was significantly higher in the Inter-ScrA condition more than the other two conditions (one-way ANOVA, F(2,57) = 6.237, p = 0.0045; Fig. 2a-c). Thus, this weaker AEP for the task-irrelevant stimulus in the Inter-SyncA and Occl-SyncA conditions provides neural evidence that the brain has reached a distinct selective-attentional state marking the flow experience. Notably, the AEP was negatively correlated with the flow index in the Inter-SyncA condition (Spearman’s Rho = −0.48, *P* = 0.03), while it was only weakly (Spearman’s Rho = −0.29, *P* = 0.22) or not correlated (Spearman’s Rho = 0.11, *P* = 0.64) with the flow index in the Occl-SyncA and the Inter-ScrA condition, respectively (Fig. 2d). These results provided an evidence that the experimental manipulations indeed produced a deep flow state in the Inter-SyncA and Occl-SyncA conditions, but not in the Inter-ScrA condition.

**Figure 2.**
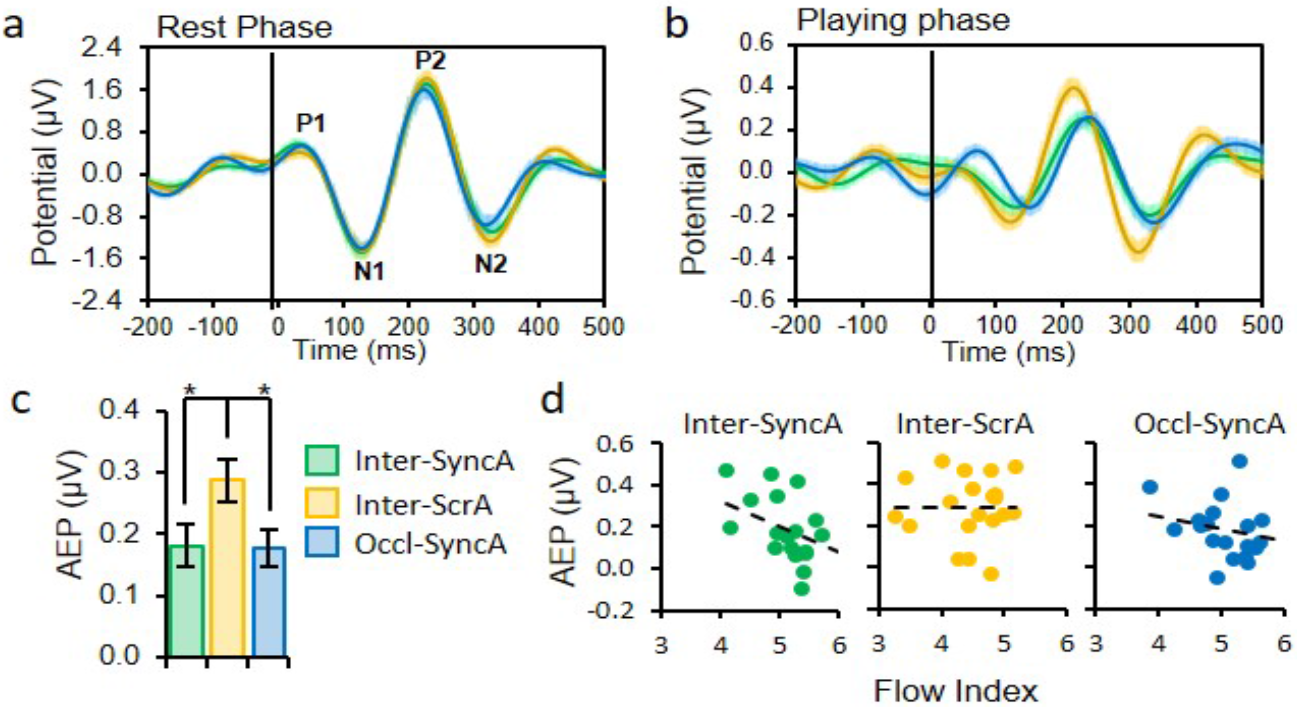
Selective-attention-based objective assessment of flow. **a, b**, The potential, pass-filtered in the theta range (3 – 7 Hz), at central channels locked to the task-irrelevant beep onsets during the resting and playing phases. **c**, The mean magnitude of the bandpass-filtered auditory-evoked potential (AEP). The non-flow condition (Inter-ScrA) showed statistically significant higher AEP than the flow conditions. One-way ANOVA with Bonferroni post-hoc test. **d**, Spearman’s correlation between AEP and flow index. AEP is negatively correlated with the flow index in the Inter-SyncA condition (Spearman’s Rho = −0.48, *P* = 0.03), showing a negative correlation trend in the Occl-SyncA condition (Spearman’s Rho = −0.29, *P* = 0.22), and no correlation in the Inter-ScrA condition (Spearman’s Rho = 0.11, *P* = 0.64). Dashed line indicates regression line. * P < 0.05. Error bars and shaded regions represent mean ± s.e.m.; n = 20.

### A unique neural signature for team flow is revealed using power spectral analysis

To detect specific neural correlates for team flow, we used power spectral analysis at various domains (Fig. S1b). First, in the frequency domain, we checked the normalized power grand-averaged across all channels for each frequency band. We found significant difference across test conditions in the alpha (8 −12 Hz; one-way ANOVA, F(2,57) = 3.5491, p = 0.0353), beta (13 – 30 Hz; one-way ANOVA, F(2,57) = 3.0756, p = 0.0539), and gamma (31 – 120 Hz; one-way ANOVA, F(2,57) = 7.7168, p = 0.0011) frequencies but not in the delta (1 – 3 Hz; one-way ANOVA, F(2,57) = 0.8067, p = 0.4514), or theta (4 – 7 Hz; one-way ANOVA, F(2,57) = 0.1693, p = 0.8447) frequency bands (Fig. S4). Second, at the topographical domain, alpha-power analysis did not show specific surface channels significantly different across conditions (data not shown). Topographical beta- and gamma-power analysis showed four channels at the left temporal area with significantly higher beta and gamma power in team flow, Inter-SyncA condition, more than the other two conditions (Fig. 3). For considerations, we used the combined beta and low-gamma (beta/gamma) band (13-50 Hz) for further analysis (please, check the Statistical analysis section). Third, at the anatomical-source domain, we performed a precise cortical source localization method that included co-registration with the individual’s structural MRI. The anatomical-source beta/gamma power showed that regions at the right and left temporal area had a significantly higher power in the Inter-SyncA condition compared to the other two conditions (Fig. 4a,b). These brain regions encompass the inferior temporal gyrus (ITS), middle temporal gyrus (MTG), and the superior temporal gyrus (STG) and sulcus (STS). Also, the beta/gamma power of these brain regions showed higher correlation tendencies with the team flow index only in the Inter-SyncA condition. For example, the Left MTG (L-MTG) showed the highest beta/gamma power correlation with the team flow index in the Inter-SyncA condition (Spearman’s Rho = 0.56, *P* = 0.006) but not in the Inter-ScrA (Spearman’s Rho = - 0.19, *P* = 0.43) or the Occl-SyncA (Spearman’s Rho = - 0.02, *P* = 0.95) conditions (Fig. 4c). The results from the power spectral analyses, at all tested domains, provided the first neural evidence that team flow is a qualitatively different brain state distinguishable from individual flow or ordinary socialization.

**Figure 3.**
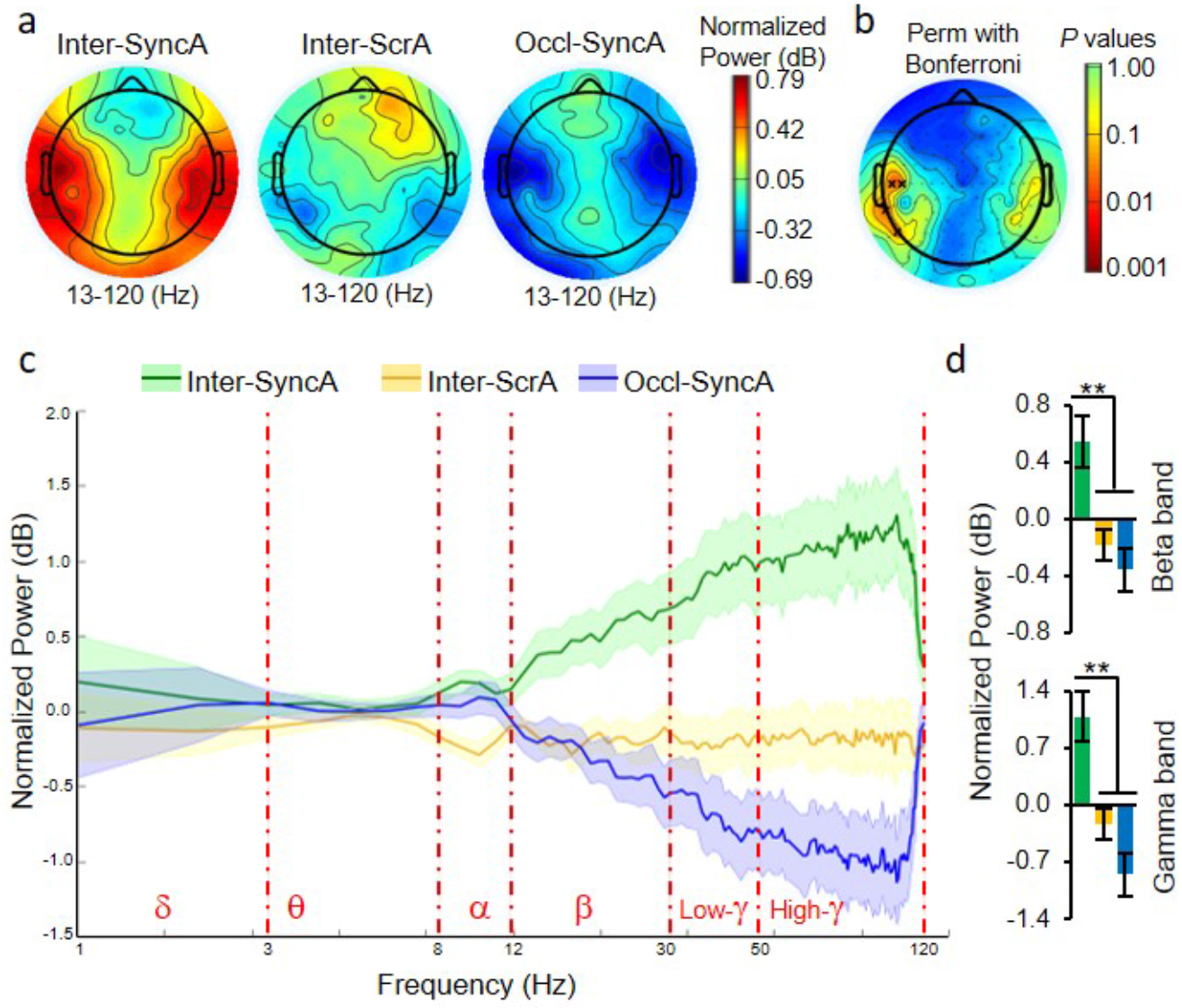
Higher beta/gamma power at the left temporal regions as a unique neural signature for team flow. **a**, The topographies of the beta and gamma frequencies (13 – 120 Hz) computed as the average over normalized power. **b**, Permutation statistical significance across conditions with Bonferroni multiple comparison corrections. The black crosses indicate channels with *P* < 0.05. **c**, The normalized power spectral analysis averaged from the four channels (D21, D22, D24, D31) in the left temporal area identified in **b**. **d**, Averaged normalized power for the beta (13 – 30 Hz) and gamma (31 – 120 Hz) frequency bands show power enhancement in the team flow (Inter-SyncA) condition. One-way ANOVA with Bonferroni post-hoc test. ** P < 0.01. Error bars represent mean ± s.e.m.; n = 20.

**Figure 4.**
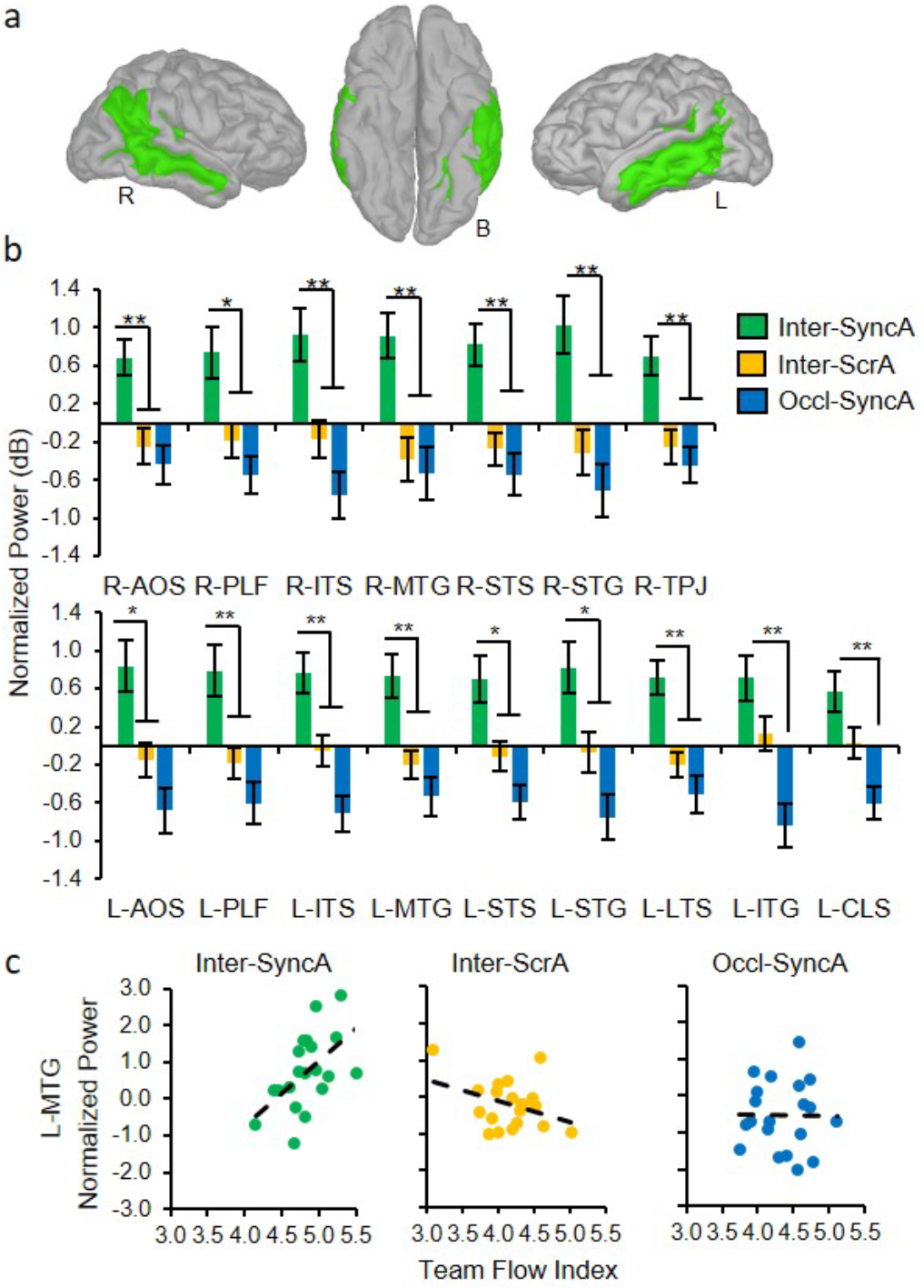
Higher beta/gamma power during team flow is anatomically localized to the middle temporal lobe and temporal-parietal regions. **a**, Brain regions, as defined by the Destrieux atlas, showing significant beta/gamma normalized power difference across conditions are highlighted in green. **b**, The average normalized power of the beta/gamma frequency band (13 - 50 Hz) at each highlighted brain region. One-way ANOVA with Bonferroni post-hoc test. **c**, Condition-specific Spearman’s correlations between beta/gamma power and team flow index at L-MTG as a representative region. Positive correlation was found in the Inter-SyncA condition (Spearman’s Rho = 0.56, *P* = 0.006), but not in the Inter-ScrA condition (Spearman’s Rho = −0.19, *P* = 0.43) or in the Occl-syncA condition (Spearman’s Rho = −0.02, *P* = 0.95). Dashed line indicates regression line. * *P* < 0.05, ** *P* < 0.01. Error bars represent mean ± s.e.m.; n = 20. B, bottom view; R, right; L, left; AOS, anterior occipital sulcus; PLF, posterior lateral fissure; ITS, inferior temporal sulcus; STS, superior temporal sulcus; STG, superior temporal gyrus; TPJ, temporal parietal junction, LTS, lateral temporal sulcus; ITG, inferior temporal gyrus; CLS, collateral and lingual sulcus.

### Unsupervised clustering analysis revealed brains areas specific to individual flow, socialization, or team flow

All the ROIs in the anatomical-domain power analysis, mentioned above, showed significant power activity specific only to team flow. This indicates that during team flow, there is no brain area still encodes individual flow or socialization. Before rushing for this conclusion, we tested an alternative hypothesis that the analysis above failed to detect weaker, but true effects, due to strict statistical threshold (check Statistical analysis section for further explanation). Since the anatomical-source localization averages source vertices based on a rigid, predefined parcellations method, we wanted to give more weight to the distribution of activity based on function rather than anatomy. Therefore, we used an activity-dependent unsupervised machine learning to cluster (cl) the source vertices based on their similarity in the beta/gamma power pattern (Fig. 5). We detected two clusters (cls), distributed over the anterior part of the frontal cortex, where the beta/gamma power was higher in the Inter-ScrA condition than the other conditions (Fig. 5; cls 1 and 2). This pattern was significant in cl 2 (one-way ANOVA, F(2,57) = 3.6125, p = 0.033), while cl 1 showed a trend (one-way ANOVA, F(2,57) = 1.5916, p = 0.2125). The suppressed activity in these clusters is specific to the flow experience, regardless of the social context, which is consistent with a neural representation of the automaticity-dimension of flow (20, 21). Also, we detected two clusters distributed mostly over the middle and inferior frontal cortex and the left occipital cortex, where the beta/gamma power was lower in the Occl-SyncA condition than the other conditions (Fig. 5; cls 3 and 4). This pattern was significant in cl 4 (one-way ANOVA, F(2,57) = 7.4841, p = 0.0013), while cl 3 showed a trend (one-way ANOVA, F(2,57) = 2.4288, p = 0.0972). The increased activity in these clusters is specific to team interactions, regardless of the flow state. The rest of the clusters were distributed mostly over the temporal, parietal, and occipital cortices, where the beta/gamma power was higher in the Inter-SyncA condition than the other conditions (Fig. 5; cls 5 - 7). This pattern was significant in all three cls: cl 5 (one-way ANOVA, F(2,57) = 11.8753, p = 0.000049), cl 6 (one-way ANOVA, F(2,57) = 9.548, p = 0.00027), and cl 7 (one-way ANOVA, F(2,57) = 6.9256, p = 0.002). The increased activity in these clusters is specific to team flow. These results indicate that even during team flow, the brain shows neural correlates of each isolated experience: individual flow and socialization. Before performing further groups interaction metric analyses, we used the activity-dependent power spectral analysis to refine the definition of brain regions to gain higher sensitivity for the neural correlates. We combined the anatomical and the functional domains mentioned above creating 14 anatomically-defined-activity-dependent groups (GPs), 7 GPs per hemisphere (Fig. S5, Table S2, and Table S3). GP7 represented the temporal brain areas, which showed significant differences across conditions (Fig. 4).

**Figure 5.**
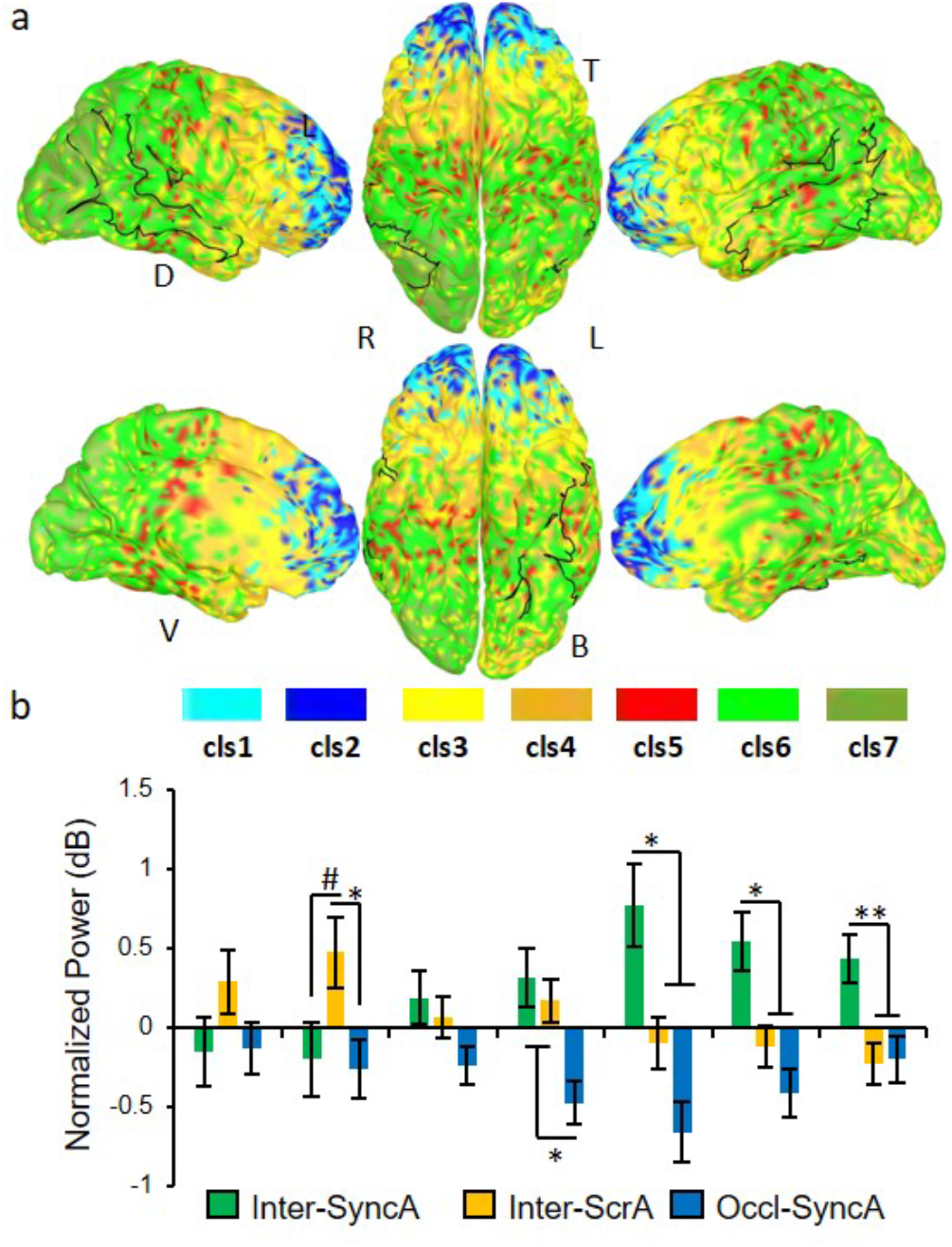
Unsupervised hierarchical vertices clustering based on beta/gamma power difference reveals flow-, social-, or team flow-related clusters. **a**, Clustered-vertices projected to a standard brain to visualize cluster localization. The black lines indicate the boundaries of the brain regions that showed significant beta/gamma normalized power across conditions as shown in Fig. 4. **b**, The cluster-averaged normalized power of the beta/gamma frequency band. One-way ANOVA with Tukey-Kramer’s post hoc test. Flow-related (cls1-2), social-related (cls3-4), or team flow-related (cls5-7) clusters are indicated in color scheme matching Fig. 5a. # P = 0.077, * P < 0.05, ** P < 0.01. Error bars represent mean ± s.e.m.; n = 20. B: Bottom; D: dorsal; L: left; R: right; T: top; V: ventral.

### The left temporal cortex receives information from brain areas encoding individual flow or socialization

The neural signature of team flow detected in the left temporal regions (L-GP7) might be the upstream neural spark that guided other brain information processes or the downstream outcome of these processes. In both cases, this neural signature will play a critical function to produce the team flow experience. Alternatively, this neural signature might be a neural byproduct passively accumulated during information processing. To disentangle these possibilities, we analyzed the causal information interactions across all the brain region groups (GPs), using three frequency-domain Granger-causality (GC) measures: the Granger-Geweke Causality (GGC), the direct directed transfer function (dDTF), and the normalized partial directed coherence (nPDC)(26). In all GC measures, the causal interaction matrix showed GP7 receives information (From) more than sending information (To) other GPs (Fig. 6a and Fig. S6a). To quantify whether a GP is a global sender or recipient of information, we calculated the global To/From ratio for each GP per condition. In the GGC measure, the logarithmic (Log) global To/From ratio for the left GP7 (L-GP7) was less than zero and significantly less than any other GP except the right GP7 (R-GP7) (Fig. 6b, two-way ANOVA, F(26,494) = 2.9768, p = 1.94e^−6^). Also, in the dDTF and nPDC measures, the global To/From ratio for the left GP7 (L-GP7) was significantly less than any other GP except the right GP7 (R-GP7) (Fig. S6b). These results indicate that the left-temporal brain regions (L-GP7) fall downstream in information causality to all other brain regions. Therefore, the detected beta/gamma power in L-GP7 might be an final information processing outcome playing a function in producing the team flow experience.

**Figure 6.**
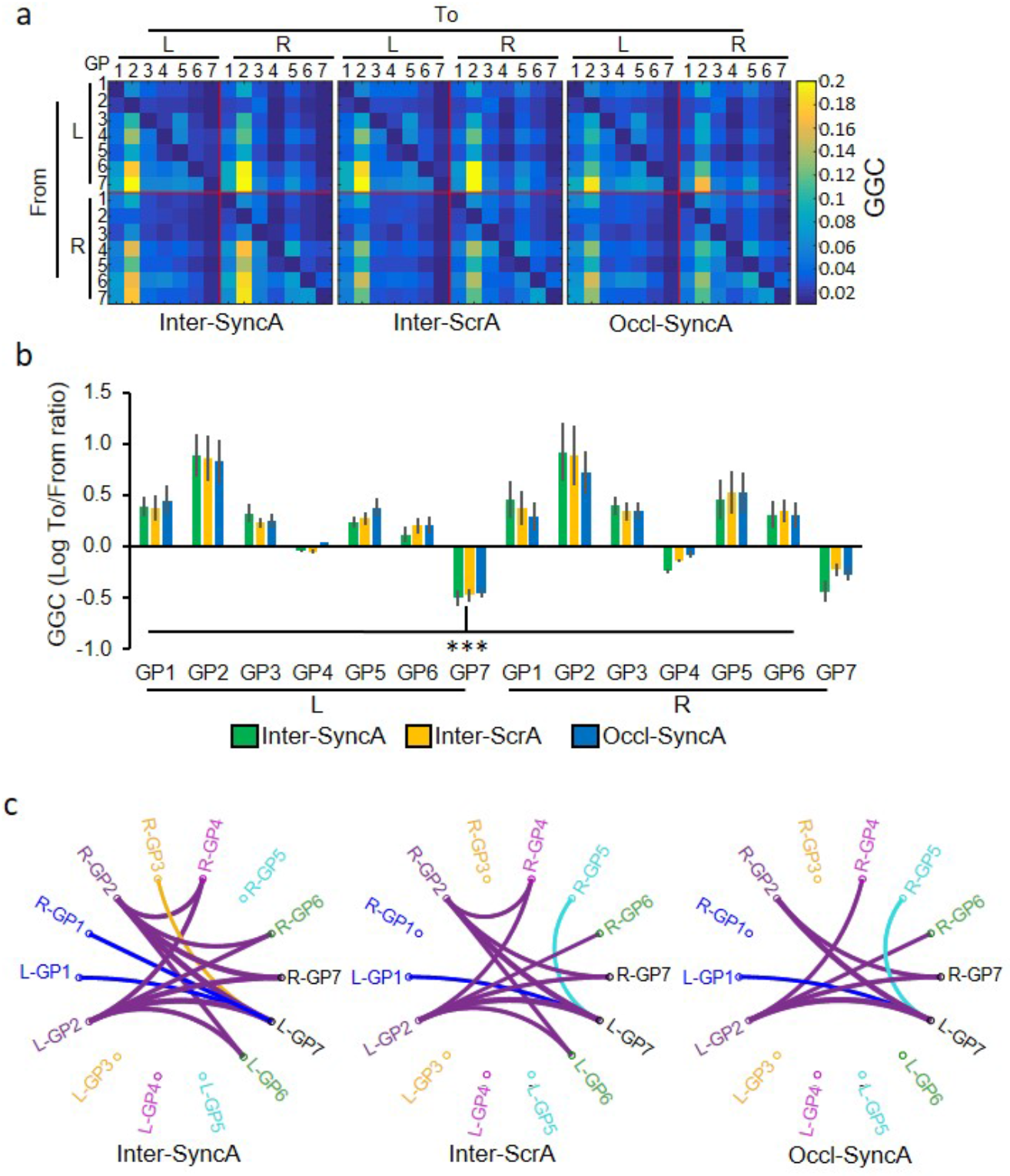
Intra-brain causality analysis showing the left temporal regions (GP7) as a downstream information recipient. **a**, The mean Granger-Geweke Causality (GGC) causal interaction matrix for the activity-dependent-anatomically-defined groups (GPs). To, indicates sending information; From, indicates receiving information; L, left hemisphere; R, right hemisphere. **b**, The mean GGC To/From ratio (log transformed) where positive values indicate global information sender to other GPs and vice versa. L-GP7 is a significant information receiver. Twoway ANOVA with Tukey-Kramer’s post hoc test. **c**, The top information senders among all GP-GP causal interactions. For each GP-GP connection, the line color matches the color of the GP name which sends the information. Notably, only in the Inter-SyncA condition, L-GP7 receives information from R-GP1 and R-GP3. *** P < 0.001. Error bars represent mean ± s.e.m.; n = 20. D-L, dorsal-left; V-L, ventral-left.

If L-GP7 contained the most downstream brain regions regarding information causality, what are the most important upstream brain regions contributing in sending information? To answer this question, for each GP-GP causal interaction, we applied a global threshold to leave only the top, approximately 10%, information senders (Fig. 6c). The top information senders are L- & R-GP1, L- & R-GP2, R-GP3, and R-GP5. Interestingly, using this threshold, in the Inter-SyncA condition, the top information senders to L-GP7 include the contralateral R-GP1 and R-GP3. On the other hand, in the Inter-ScrA and Occl-SyncA conditions showed a different causality pattern where the top information senders to L-GP7 include the contralateral R-GP5. Collectively, these results indicate that the unique beta/gamma power detected in the left temporal regions (L-GP7) is an outcome of information processing that happened earlier in time and the sources of these information include brain areas that encodes individual flow (GP1) and socialization (GP3).

### The left temporal cortex is involved in integrated information at both the intra- and inter-brain levels

In the Inter-SyncA condition, L-GP7 received information from R-GP1, where the beta/gamma activity is related to individual flow, and from R-GP3 where the beta/gamma activity is related to social interaction (Fig. 5b and 6c). On the phenomenological level, it is hard to decompose the team flow experience into two isolated components; flow and socialization. Therefore, a possible outcome of such causality is an effect on the integration of information from the brain regions coding each isolated experience. To test this hypothesis, we used the integrated information theory (27, 28). We calculated the normalized Integrated Information value (Norm II) as a metric for the integrated information. In both intra-brain and inter-brains calculations, there was a general tendency for Norm II to be higher in the Inter-SyncA condition than the other conditions (Fig. 7a). When we averaged the Norm II across all GP-GP connections (Global Norm II), the Inter-SyncA showed significant higher inter-brains Global Norm II than other conditions (one-way ANOVA, F(2,57) = 15.0516, p = 5.64e^−6^), while showing similar trend at the intra-brain level (one-way ANOVA, F(2,57) = 3.6251, p = 0.033) (Fig. 7b). Next, we checked all GP-GP connections for a significant Nom II at Inter-SyncA condition compared to other conditions (three-way ANOVA, condition x GP1 interaction for intra-brain: F(26,10133) = 4.7622, p = 1.35e^−14^, and for inter-brain: F(26,10959) = 3.676, p = 7.76e^−10^). Among all GP-GP connections, significant connections were detected only at the left hemisphere as detailed in Fig. 7c. These connections formed an intra-brain L-GP3-GP4-GP5-GP7 subnetwork and an inter-brains L-GP7-to-L-GP7 link that showed significant higher Norm II in the Inter-SyncA (Fig. 7c). These results indicate that during team flow, the team members exhibited higher information integration not only within each player’s brain but also between their brains. More specifically, L-GP7 was the only group of brain regions that showed significantly higher inter-brain integrated information during team flow.

**Figure 7.**
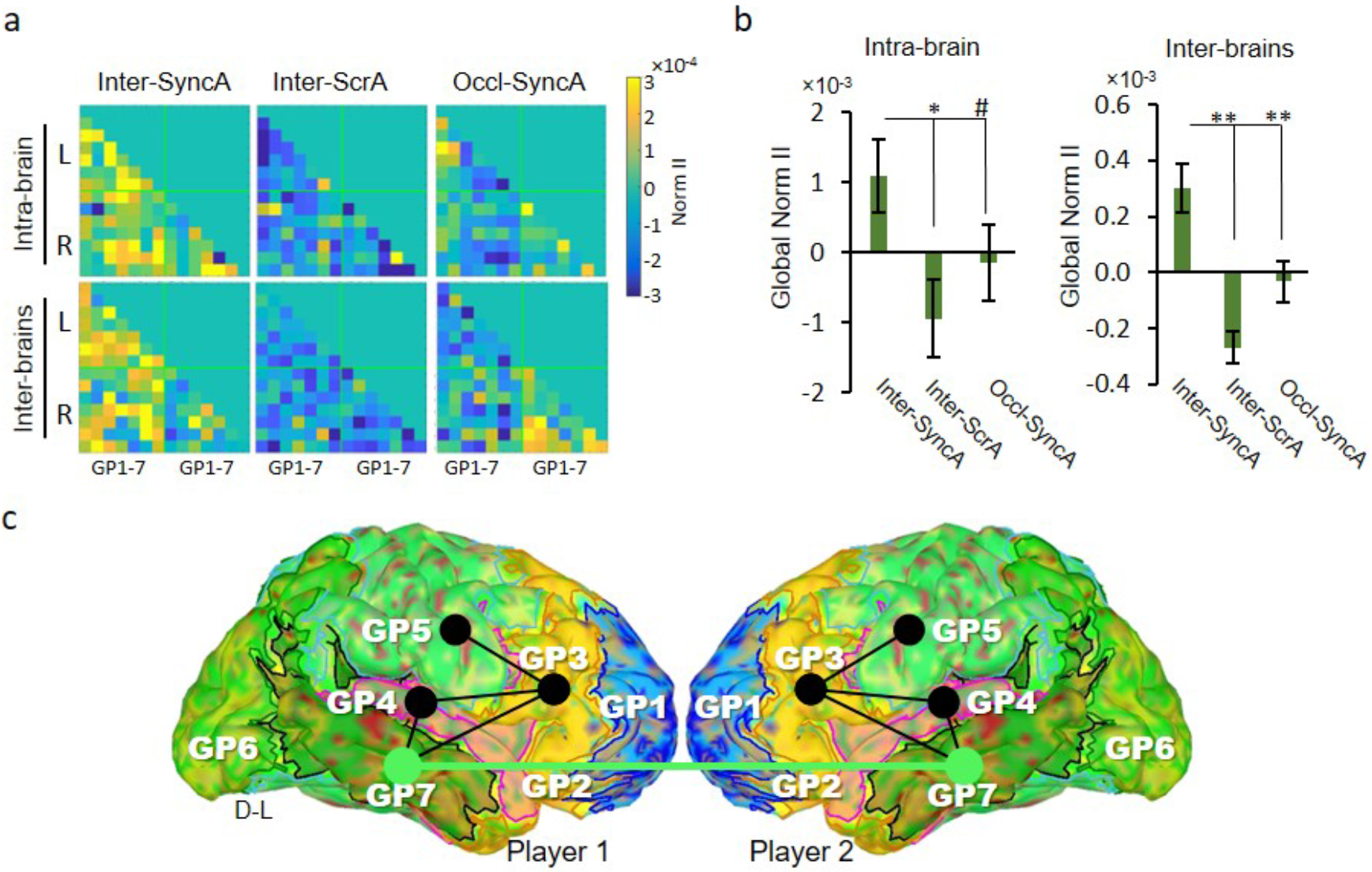
Team flow showing enhanced integrated information globally and specifically at the left temporal regions (GP7). **a**, The mean normalized Integrated Information value (Norm II) connectivity matrix for the activity-dependent-anatomically-defined groups (GP1-7). Normalized Integrated Information is presented as difference from the average Integrated Information for each GP-GP connection across conditions. **b**, The mean Global Norm II averaged across all GP-GP connections showing significantly higher inter-brains (left panel) and intra-brain (right panel) during inter-SyncA condition. One-way ANOVA with Bonferroni post hoc test. **c**, GP-GP connections that shows significant (P < 0.05) Norm II in the inter-SyncA condition compared to other conditions. Three-way ANOVA with Bonferroni post hoc test. Black lines indicate intra-brain and green line indicates inter-brains GP-GP connections. D-L, dorsal-left. * p < 0.05, ** p < 0.01, # p < 0.1. Error bars represent mean ± s.e.m.; n = 20.

### Team flow is associated with higher intra- and inter-brain neural synchrony

The enhanced inter-brain integrated information might concur with enhanced neural synchrony between the team’s brain regions. To check for this hypothesis, we calculated the inter-brains Normalized Phase-Locking Values (Norm PLV) across all the GP-GP connections for each condition (Fig. 8a). The results showed a general tendency for Norm PLV to be higher in the Inter-SyncA condition than other conditions at both the intra- and inter-brains level. The inter-brain Norm PLV calculated using a randomly shuffled pairs does not seem to show any difference across conditions. To quantify this effect, we averaged the Norm PLV for all GP-GP connections (Global Norm PLV). The Inter-SyncA showed a significantly higher Global Norm PLV than other conditions only in the actual paired participants but not in randomized pairs (two-way ANOVA, F(2,114) = 3.416, p = 0.0362) (Fig. 8b). Collectively, these results indicate that during team flow, the team members exhibited higher integration and neural synchrony between their brains. This enhancement in information integration and neural synchrony is consistent with the phenomenological experience during team flow. Indeed, it might be the neurocognitive basis for the superior subjective experience of team flow.

**Figure 8.**
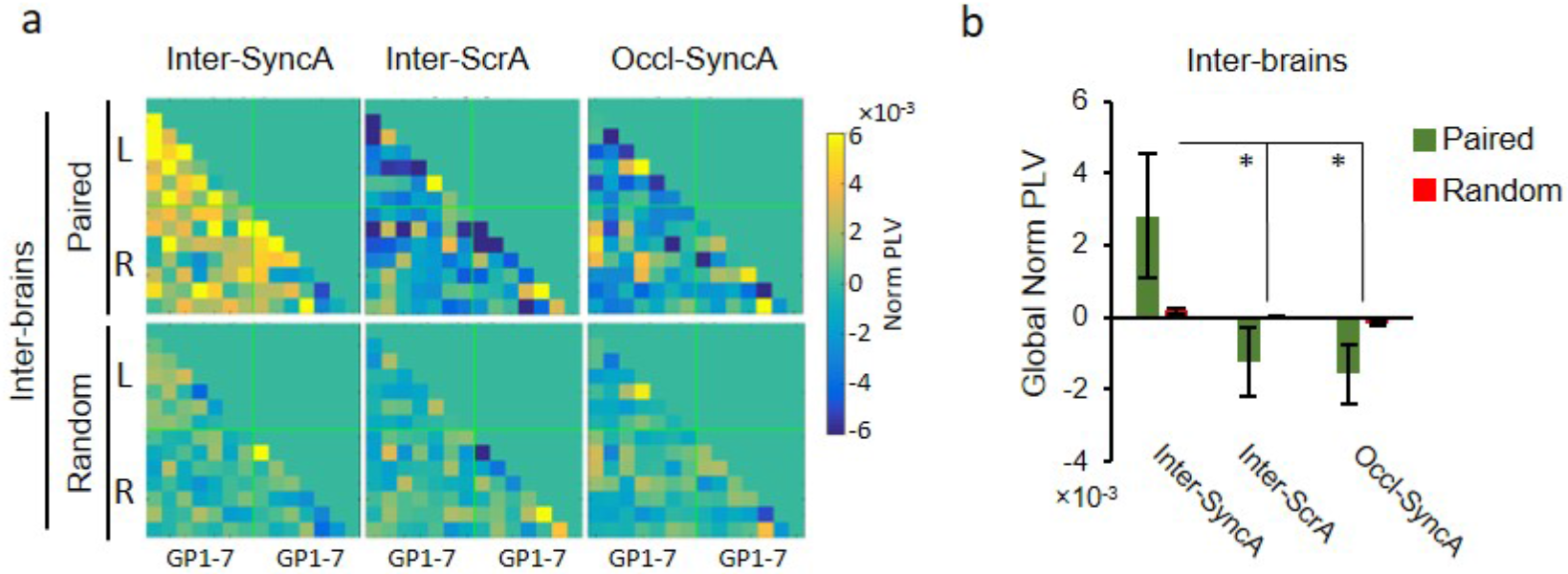
Phase-locking value showing a global enhanced inter-brain synchrony during team flow. **a**, The mean phase-locking value (PLV) connectivity matrix for the activity-dependent-anatomically-defined groups (GP1-7). Normalized PLV is presented as difference from the average PLV for each GP-GP connection across conditions. Paired, indicates the actual experimental pair; Random, indicates randomly selected pairs. **b**, The mean Global normalized PLV averaged across all GP-GP connections showing significantly higher inter-brains during Inter-SyncA condition. Two-way ANOVA for inter-brain with Bonferroni post hoc test. * p < 0.05. Error bars represent mean ± s.e.m.; n = 20.

## DISCUSSION

### Effective parametric tools to measure the depth of the flow experience

So far, the flow state was identified in the literature using only subjective questionnaires, which we have reproduced in this report (Fig. 1 and Fig. S2) (20–23). Although psychometric ratings provided some evidence that the participants reached some sort of the flow experience, it did not give us reliable clues about the depth of the flow state (24). Using the task-irrelevant AEP, we could confirm that the task used in this report could attain enough depth of flow experience to alter one of the most critical flow dimensions: the attenuated consciousness to external stimulus. Indeed, the flow experience is hypothesized to be either an abrupt discrete zone or a gradual continuum (21, 24). Solving this ambiguity would be a significant advance in understanding how the flow experience develops and operates. Our newly developed method for measuring the flow depth can be a useful parametric tool to help to answer this question.

### The unique neural correlates of team flow and it generality

The most prominent neural correlate for the team flow state identified in this study is the higher beta/gamma power in the temporal regions (GP7) as shown in Figs. 3, 4 and 5. A critical question is whether this unique team flow signature is only related to the particular task employed in this report i.e., the music rhythm game or representing a general correlate to team flow regardless of the task. Beta and gamma oscillations are involved in several cognitive functions, including attention, memory, and awareness, with evidence of abnormalities in brain disorders (29). In general, these functions are consistent with higher team interactions, and with enhancement in many flow dimensions. Moreover, our data agree with the neural activities detected in other reports, studying individual flow or social interaction, that used completely different tasks. For example, in GP1 (the anterior part of the frontal cortex), the beta/gamma power was lower in the flow conditions compared to the non-flow conditions (Fig. 5 and Fig. S5). These data agree with the reduced activities in the medial PFC in an arithmetic task (21, 23). Also, in GP3 (that includes the IFG), the beta/gamma power was higher in the team conditions compared to the non-team conditions (Fig. 5 and Fig. S5). These data agree with the involvement of the IFG in social interaction in a plethora of different tasks (14, 30). These oscillations and brain regions are seemingly not constrained to a specific task but specific functions supporting the generality of our conclusions. Still, the ultimate test is to reproduce the team flow experience, and its neural correlates in other situations and tasks, which goes beyond this report.

### A suggested neural model for team flow

As mentioned earlier, team flow might require a unique interaction between the brain regions involved in individual flow and socialization. As reported, the PFC are activated during social interaction but suppressed during individual flow experience (14, 15, 24). During team flow, the PFC showed both signals. The anterior part of the PFC (GP1) showed lower activity related to the effortless information processing during the flow experience. On the other hand, the posterior part (GP3) showed high activity associated with more social information processing. These observations argue that the neural correlates of team flow are unique in terms of the interaction of the different neural networks, not just a simple addition of known functional modules of individual flow and socialization.

Our data also show that L-GP7, the left temporal cortex, falls downstream to other brain areas and receives information from brain areas encoding individual flow, GP1, and from areas encoding social interaction, GP3. Also, L-GP7 was the only significant region showing higher integrated information during team flow at both the intra- and inter-brain levels. Both of the above results imply that the left temporal cortex function as a region that collects flow-related and social-related information and integrate them during the team flow experience. Previous reports also suggest an integration function for the temporal cortex in different contexts. For example, the middle and inferior temporal gyrus, regions included in GP7, have been reported to play a role in cognitive-affective integration in schizophrenia (31). Also, superior temporal sulcus, anther region within GP7 and has been extensively reported in social information processing, played a role in integrating multiple social networks (15). Thus, our results and past reports fall in line to suggest a neural model during team flow where the left temporal cortex is involved in integrating the flow and social information to serve the team flow experience, as depicted in Fig. S7.

### Team flow as a bonafide inter-brain state

Recent social neuroscience studies have reported interactions between the brains of a group of members measured as inter-brain synchronization (e.g., phase synchrony). This synchrony can be enhanced during intense socialization, body or speech coordination, music production, dancers, student-teacher interactions in classrooms, touch-mediated pain reduction, creativity in cooperative tasks, and even in socially interacting bats (32–41). Hence, it can be a metric for more effective group interactions. Similarly, integrated information, which measures the amount of information generated by the system as a whole relative to the sum of its parts, is another metric of group interaction (28). Integrated information between the brain can predict effective group interaction and complexity; thus, it can be linked to collective intelligence (42). Crucial data in our report is the significantly higher inter-brain information integration and neural synchrony during team flow (Fig. 7 and 8). Based on both metrics, our data indicate that team flow creates a hyper-cognitive state between the team members.

One might argue that these results are the outcome of the pair of participants sharing the same sensorimotor information. Hence, the synchronized neural substrates are related to a lower-level task-related cognitive function. This interpretation is not valid as the pair of participants shared the same sensorimotor information in all the three experimental conditions. We were careful in our experimental design to refute such a conclusion by keeping almost all the visual, auditory, and motor information constant across conditions (Table S1). We only used minimum manipulations to create the proper subjective experience. Even with such minimum manipulations, our study kept a real-life dynamic interaction between participants (9, 34, 43). These dynamic interactions were essential and sufficient to induce the team flow experience and its unique brain state.

In this context, whereas the integrated information is typically considered a quantitative measure of consciousness (28), there is considerable debate among researchers about whether integrated information is a sufficient measure of consciousness (44). Hence, our findings of the high value of integrated information do not necessarily indicate a modified form of consciousness, for instance, “team consciousness.” Nonetheless, its consistency with neural synchrony (PLV) certainly raises intriguing and empirical questions related to inter-brains synchrony, information integration, and altered state of consciousness.

## MATERIALS AND METHODS

### Participants

In the main experiment, 15 participants (5 males; age: 18-35 years) attended forming 10 pairs (3 male pairs) where 5 participants (1 male) were paired twice. Written informed consent was acquired from all participants. All the procedures were approved by the Institutional Review Board of the California Institute of Technology.

### Task

We used a commercial music rhythm game called “O2JAM U” (version 1.6.0.11, MOMO Co., South Korea). We used the 4-lane mode during the main experiment where a pair of participants played with each participant responsible for two adjacent lanes. Task, manipulations, screening and main experimental sequence are detailed in the SI.

After playing each song (trial), the game displays a performance report on the screen, including a final numerical score, the total number of cues, and the number of missed cues. The performance report of each trial was hidden from the participants until they finish answering their subjective experience questionnaire. The percentage of the missed cues per the total number of cues was used as a metric for the performance of each pair of participants.

### Task-irrelevant stimulus

Task-irrelevant auditory stimulus (beep sound) were pseudo-randomly presented to probe the strength of the participants’ selective-attention to the game and was used as an objective measure of flow. We presented beep trains played at 5 Hz for 1 second (i.e. each train consisted of 5 beeps). Each beep was at 500 Hz and lasted for 10 milliseconds. The beep trains simulated the sound of someone knocking on a door to make the stimulus as natural as possible. The interval between the beeps train varied from 4 - 8 seconds. The beeps were generated by Matlab 2012 (The MathWorks, Inc., Natick, Massachusetts, United States) and delivered through another pair of speakers placed equidistant from the iPad.

### Anatomical MRI acquisition

To increase the accuracy of source estimation for cortical activity, individual head anatomy from each participant, who passed the screening and agreed to participate in the main experiment, was acquired with magnetic resonance imaging (MRI). A 3 Tesla Siemens Trio (Erlangen, Germany) scanner and standard radio frequency coil was used for all the MR scanning. High resolution structural images were collected using a standard MPRAGE pulse sequence, providing full brain T1-weighted 3-dimensional structural images.

### Psychometric ratings and calculation of experience indices

From subjective reports, we calculated the flow, team, and team flow indices, by calculating the arithmetic mean of the ratings for each trial, to estimate subjective experience for individual flow, positive team interaction and team flow, respectively (Fig. S2). For assessing flow experience, questions related to the skill-demand balance (Q1 and Q2), feeling in control (Q3), automaticity (Q4), enjoyment (Q5), and time perception (Q6) dimensions of flow were used(2). For assessing team interaction, questions related to awareness of partner (Q7), teamwork (Q8), and coordination (Q9) dimensions of positive team interaction were used. Questions assessing competition (Q10) and distraction (Q11) were asked to confirm the absence of negative team interactions and were not included in any index. The team flow index was calculated by averaging the flow and team indices.

### Hyperscanning EEG recording and preprocessing

Electroencephalogram (EEG) was recorded simultaneously from both participants using a dual BioSemi ActiveTwo system (BioSemi Inc., Amsterdam, The Netherlands). Each participant wore a cap holding 128 scalp Ag/AgCl electrodes. Signals were amplified by two daisy-chained ActiveTwo AD boxes where one AD box was connected to the control PC and worked as a master controlling the other AD box to ensure synchronization. Electrode impedance was kept below 10 kΩ. For each cap, an active Common Mode Sense (CMS) electrode and a passive Driven Right Leg (DRL) electrode positioned near vertex served as the ground electrodes. EEG signals were recorded at a sampling rate of 2048 Hz (later down-sampled to 256 Hz). During recording, the A1 electrode, or A2 electrode in 3 participants, served as a reference. In the ABC layout (a Biosemi designed equiradial system), these electrodes overlap with the Cz location of the international 10–20 system. Signals were recorded and saved using ActiView/LabView software (version 8.04, BioSemi Inc., Amsterdam, The Netherlands) installed on the control PC. Another master PC was used to generate the task-irrelevant beep sound and to send signals to the EEG data receiver marking the onset of each beep train (event triggers). The event triggers were used to align the EEG data with the resting and the playing phases by using a real-time projection of the top-view video recording to the control PC. The experimenter confirmed that all the onsets of the beep trains happened during the resting or the playing phase periods.

To analyze the auditory-evoked potentiation (AEP), EEG data were epoched −0.5 sec to 1 sec (1.5 sec total) flanking the beep train onsets (AEP epochs). To analyze the neural correlates (NC) of game play experience, EEG data were epoched 2 sec to 5 sec (3 secs total) after the beeps train onset (NC epochs; Fig. S1b). EEG data was high-pass filtered at 0.5 Hz, re-referenced to the average of all channels and artifact corrected using automatic independent component analysis (ICA) rejection using the FASTER toolbox (45). Bad channels showing line noise noted during recording sessions were rejected and interpolated during the FASTER preprocessing.

### Auditory evoked potential analysis

To select the channels maximally responsive to the task-irrelevant auditory stimuli, we analyzed the AEP epochs during the resting phase. We calculated the event-related spectral perturbation (ERSP) and the inter-epochs coherence (IEC) using EEGLAB toolbox (version 14.1.1) (46). Both ERSP and IEC showed changes in theta activity (3 – 7 Hz) at 100 – 350 ms post-onset with a peak increase at 150 – 250 ms post-onset (Fig. S3a,b). Topographical analysis in the theta band showed a strong positive activity in the 14 central channels from 200 - 260 ms post-onset (Fig. S3c). The frequency, time and topographical frames of our AEP were consistent with previous reports (47, 48). For each trial, we used IEC in the theta band during the resting phase to select channels showing stable AEP. IEC was averaged across the 14 central channels and channels showing IEC lower than one standard error below the mean were excluded from further AEP analysis for that trial. We then analyzed theta power from −200 ms to 500 ms flanking the beep train onsets during the resting and the playing phase (Fig. 2a,b). AEP peak amplitude was calculated according to the method described by a previous simulation report showing that event-related potential measured based on the mean amplitude surrounding the group latency is the most robust against background noise (49). Therefore, we first calculated the N1, P2, and N2 peak latencies averaged across all conditions during the resting phase (Fig. 2a). Second, the individual N1, P2, and N2 mean peak amplitudes ± 40 ms surrounding the calculated peak latencies were obtained during the playing phase (Fig. 2b). This results in the following time windows: N1 (110 – 150 ms), P1 (210 – 250 ms), and N2 (310 – 350 ms). The theta power from the selected channels averaged across these time windows was used as AEP (Fig. 2c,d).

### Anatomically-defined source estimations

FreeSurfer (50) was used for automatic segmentation and reconstruction of the MRI images. MRI images were used to compute each individualized head model using boundary element model (BEM) implemented in OpenMEEG within BrainStorm software package (version 3.4) using the default parameters (51, 52). MRI registration with EEG electrodes head positions was aligned with each participant’s BEM model and sources were computed (version 2018) using BrainStorm for each NC epoch in the playing phase. Maps of cortical activity density were obtained across the BEM mesh using the distributed minimum-norm estimate (MNE) method; with constrained dipole orientations and no baseline noise correction. For cortical region-based analysis, brain regions were defined according to the anatomical parcellation of the Destrieux atlas as implemented in FreeSurfer and available in BrainStorm (53). The time series of source activities from the 15002 vertices and the averaged activity of the predefined 148 regions of interest (ROIs) were exported for further analysis.

### Power spectrum analysis

Power spectral density (PSD) estimate was calculated using Welch’s overlapped segment averaging estimator as implemented in the MATLAB 2016a signal processing toolbox within the EEGLAB toolbox using default parameters (46, 54). The normalized PSD was calculated for each NC epoch then averaged within each trial yielding trial PSD data at each of the 128 channels, the 148 brain region sources, and the 15002 mesh vertex sources. For each song played, the individual’s mean PSD across the three conditions was calculated. The normalized power was calculated through subtracting the individual’s mean PSD from the PSD at each condition. The normalized power was averaged within the following frequency bands: delta (1 – 3 Hz), theta (4 – 7 Hz), alpha (8 −12 Hz), beta (13 – 30 Hz), gamma (31 – 120 Hz), and lower gamma (31 – 50 Hz). The normalized power for the 128 channel data and the permutation statistics with Bonferroni multiple comparison correction was projected to topographical maps using EEGLAB toolbox. As detecting high-gamma power (> 50 Hz) using noninvasive EEG might be prone to artifacts (55), we only considered the combined beta and low-gamma (beta/gamma) band (13-50 Hz) for further analysis. We used one-way ANOVA across condition for determining the significance in each anatomical-source beta/gamma power effect. We set the significance threshold to p < 0.00034 (i.e. 0.05 / 148 ROIs) to correct for multiple comparison.

### Unsupervised clustering analysis

We clustered the 15002 mesh vertex sources based on their beta-gamma power. We used scikit-learn, a Python machine learning library, and implemented the unsupervised agglomerative clustering approach (56). Agglomerative clustering uses a bottom-up hierarchical approach where vertices are progressively linked together into clusters based on their feature similarity. We used 3 features for clusters which are the grand averaged beta-gamma normalized power at each of the 3 conditions. We used the Euclidean distance as a similarity measure, and the complete linkage criteria which minimizes the maximum distance between observations of pairs of clusters. We have tried setting the number of clusters into 3 – 8 clusters. We selected the minimum number of clusters, 7 in our case, that would extract statistically significant flow- and team-related clusters. When we set the number of clusters to 3 up to 6 clusters, we get similar trend of results but did not pass significance using one-way ANOVA.

### Activity-dependent anatomically-defined grouping of ROIs

First, for each anatomical-defined ROI, we calculated the cluster composition as the percentage of the flow-related clusters (cls1-2), team-related clusters (cls3-4), and team flow-related clusters (cls5-7). Second, we checked if the anatomically-defined ROIs can be spatially subdivided into smaller ROIs with clear tendencies for a certain activity-dependent cluster composition (Fig. S5a,b). This check was done by calculating a cumulative cluster composition curve to define a threshold for subdividing the ROIs (Fig. S5b). We presented superior frontal cortex as an example of the subdivided ROIs (Fig. S5a). Next, we grouped anatomically-defined ROIs or their subdivisions into 7 groups (GPs) per hemisphere (GP1-7) based on the major activity-dependent cluster composition (Fig. S5c). For each of the 14 GPs, the activity-dependent cluster composition is summarized in Table S2 and the anatomical composition is summarized in Table S3. Anatomically-defined ROIs that showed significant beta-gamma normalized power across conditions, as shown in Fig. 4, were grouped as GP7 regardless of their composition. The time series from all the 15002 vertices were averaged based on the new 14 GPs and hence, reduced into 14 time series for each trial per participant.

### Intra-brain causal interactions analysis

We used the Source Information Flow Toolbox (SIFT) to fit an adaptive multi-variate autoregressive (AMVAR) model for the 14 GPs activities for each subject’s trial using the Vieira–Morf algorithm (57). We fitted the NC epoch with a sliding window length of 500 ms and a step size of 25 ms (26). Model order was selected by minimizing the Akaike Information criterion. We validated each fitted model using tests included in SIFT for consistency, stability, and whiteness of residuals. To estimate causal interactions, we used three directed modelbased linear frequency-domain Granger-causality measures (26). These measures are the normalized partial directed coherence (nPDC)(58), the direct directed transfer function (dDTF)(59), and the Granger-Geweke Causality (GGC)(60, 61). For each connectivity measure, we averaged across trials for each participant per condition, then averaged across the NC epoch time interval (3 secs) and across the beta-gamma (13 – 50 Hz) frequency. Finally, to quantify the degree by which a GP sends or receives information, we calculated the ratio of sending (To) divided by receiving (From) for each GP-GP interaction then average these ratios for each GP per condition per participant (To/From ratio). Two-way ANOVA was used as statistical test. To calculate the information senders for GP-GP causal interactions, we used the Log To/From GGC ratio for each GP-GP connection. Top information senders were calculated by setting a threshold where the p-value of 0.064. The GP-GP connection above this threshold were represented on a circular graph.

### Integrated Information analysis

Integrated Information (II) was used as a measure of inter-GP bidirectional causal interaction. For every pair of time courses of the GPs activities, within and between participants, we operationalized the “state” of the pair of GPs by discretizing time-samples into binary values. To roughly match the frequency range of 13-50 Hz, we first down sampled the GPs activities to give timesteps of 12.8, 17.1, 25.6Hz or 51.2Hz (that is, a time step of 19.5, 39.1, 58.6, or 78.1 ms). Using the down sampled GPs activities, we then converted each pair of consecutive time samples to “on” if the GPs activity’s voltage was increasing over two time steps and “off” otherwise. Using the time series of binarized states, we computed the probabilities of each state transitioning into each other state, constructing a transition probability matrix (TPM) which describes the evolution of the pair of GPs activities across time. To ensure accuracy of transition probabilities, we computed these across all trials. As lower time resolutions give fewer observations with which to compute the probabilities, we repeated the down sampling for each possible “start” (i.e. for each time-sample in the first time-bin) and used all transitions from all shifted-down sampled time series to build the TPM. We then submitted the TPM to PyPhi (1.2.0)(62), which then constructs a minimally reducible version of the TPM, assuming independence of GPs activities, and compares the original TPM to the minimally reducible version to compute Integrated Information (28, 62). For each actual pair, we calculated normalized Integrated Information value by subtracting the absolute value from the average across all conditions for each GP-GP connection. Three-way ANOVA (condition × GP1 x GP2) was used as statistical test for normalized Integrated Information at each GP-GP connection. The Global normalized Integrated Information was calculated through averaging normalized Integrated Information values across all possible GP-GP connections. One-way ANOVA was used as statistical test for Global normalized Integrated Information.

### Phase synchrony analysis

The phase-locking value (PLV), or inter-site phase clustering (ISPC), was used as an index of neural synchrony. The distribution of the phase angle differences between sources was generated at each time point (within the NC epoch 3 sec window) then averaged over (ISPC-trial) (63, 64). ISPC-trial was calculated at each frequency and then averaged across the frequency band of 13 – 50 Hz. For each condition, we calculated the ISPC-trial between all sources for the actual pairs or for each of 10 randomly assigned pairs. For each actual or random pair, we calculated normalized PLV value by subtracting the PLV value from the average across all conditions for each GP-GP connection. The Global normalized PLV was calculated through averaging normalized PLV values across all possible GP-GP connections. ANOVA was used as statistical test.

### Statistical analysis

All statistics were done using the Statistics and Machine Learning Toolbox within MATLAB 2016a. Authors acknowledge that most of the analyses done in this study were exploratory. In this section, we are giving a parameter justification for each analysis based on the rationale for doing the analysis.

- Screening process: the screening process is inevitable in this study to attain reasonable team flow behavioral response. In the first screening process, we needed to assure that participants signed to this study have enough skill to fall into the individual flow state. In the second screening process, we needed to match participants based on their skill and song preference. We assumed that this screening would maximize the chances of finding pairs of participants who can reach the team flow state. This quicker and cheaper behavioral-based screening process was necessary before committing to more lengthy and expensive neuroimaging process.

- Sample size: The final number of participants was mainly constrained by availability after the screening process. We strived to keep the final number of participations similar to the samples sizes reported in similar publications (34). Note: during the main experiment, the data collection process for one male pair of participants was interrupted due to a technical error and the collected data was excluded from data analysis.

- Trial numbers: we limited the number of trials to 6 per condition to avoid fatigue which might have compromised the possibility of falling into the flow state in later trials. Note that the total experimental time for each pair was approximately 3-4 hours including co-registration, EEG cap set up, the main experiment and clean up. Note: for one pair, we could only collect 5 trials per condition due to time constraints. For another pair, one of the trials contained excessive noise and hence, we excluded this trial and all corresponding trials in the other conditions.

- Units of analysis: for all analyses conducted in this study, the unit of analysis was participation i.e. n = 20. The only exception was the performance analysis where the unit of analysis was the final score for the pair i.e. n = 10.

- Control conditions: we paid much attention in the design of our manipulations to control for all possible sources of neural variability other than what can be attributed to the desired behavioral state or condition. Please, check Table S1 for details. Data collection were not performed blind to the conditions of the experiment. Experimental blinding was not possible due to the overt and obvious nature of the experimental setup for each manipulation. Data in all conditions were subjected to identical analysis algorithms.

- Spectral analysis: The power spectral analysis was completely exploratory. For the topographical domain power spectral analysis, we used permutation statistics with Bonferroni multiple comparison correction. Authors did explore the anatomical-source domain power spectral analysis and cluster analysis using beta (13-30 Hz), gamma (31 – 120 Hz), beta/full-gamma (13-120 Hz), and beta/low-gamma (13-50 Hz) bands. All bands showed similar trends with the beta band showing the least and the beta/full-gamma band the most significant effect across conditions. However, there are some limitations to the capability of EEG to accurately detect high-gamma (> 50 Hz) power. Therefore, we decided to use combined beta/low-gamma (13-50 Hz) for further analysis.

For the anatomically-domain power analysis, we restricted our anatomical atlas definition to the Destrieux atlas. The search for an ROI showing significant effect across conditions were exploratory and using one-way ANOVA across conditions. Therefore, we set the significance threshold to p < 0.00034 (i.e. 0.05 / 148 ROI) to correct for multiple comparison. All these ROIs showed higher beta/gamma power in the Inter-SyncA condition compared to other conditions as presented in Fig. 4.

- Cluster-analysis: as we set strict significance threshold in the anatomically-domain power analysis, we might have committed a type II error by ignoring true effects of other trends. The strict significance threshold was set by the number and rigid anatomical definitions of the atlas. Therefore, we used this analysis to check for the possibility of weaker but still significant trends. Hence, we increased the number of clusters just enough to detect these weaker trends. We needed to detect these weaker trends to answer the question: whether the brain still shows neural correlates of individual flow and socialization during team flow. This also justify the need for redefining the brain regions as mentioned in the Activity-dependent anatomically-defined grouping of ROIs section. Note that we have tried several clustering algorithms and parameters, all gave similar results.

- Causal interaction, IIT, and phase synchrony analyses: these analyses were done on the 14 GPs defined in the Activity-dependent anatomically-defined grouping of ROIs section. Note that all these analyses are orthogonal to the basis by which these GPs are defined which is beta/gamma power.

## Acknowledgments

We thank Dr. Chalres Yokoyama (University of Tokyo, Japan), Dr. Simone Shamay-Tsoory (University of Haifa, Israel), Dr. Katsumi Watanabe (Waseda University, Japan), and Dr. Makio Kashino (NTT communications, Japan) for their comprehensive comments on the manuscript. We thank Naomi Shroff-Mehta (Scripps College, CA), Salma Elnagar (University of Cambridge, UK), and Shota Yasunaga (Pitzer College, CA) for help with data collection and analysis. We thank Wenqi Yan (Monash University, Australia) for preliminary data analysis with integrated information. This work is supported by the program for promoting the enhancement of research universities funded to Toyohashi University of Technology to M.S. and S.N.; and the Japan Science and Technology (JST)-CREST Grant Number JPMJCR14E4 to S.S. M.C. is supported by the University of Hong Kong postgraduate scholarship program. C.T. is supported by the University of Hong Kong General Research Fund. N.T. is supported by Australian Research Council Discovery Projects (DP180104128 and DP180100396). A.L. is supported by an Australian Government Research Training Program (RTP) Scholarship.

## Author Contributions

M.S., M.C., and S.S. designed the experiments. M.S., M.C., and D.W. performed the experiments. M.S., M.C., S.S., A.L. and N.T. analyzed the data. M.S., S.S., M.C., N.T., D.W., S.N., and C.T. wrote the manuscript.

## REFERENCES

1. M. Csikszentmihalyi, Beyond boredom and anxiety, Jossey-Bass behavioral science series (Jossey-Bass Publishers, San Francisco, ed. 1st, 1975), pp. xviii, 231 p.

2. J. Nakamura, M. Csikszentmihalyi, The concept of flow. Handbook of positive psychology., Handbook of positive psychology. (Oxford University Press, 2002).

3. L. Harmat, Flow experience : empirical research and applications (Springer Berlin Heidelberg, New York, NY, 2016), pp. pages cm.

4. M. Csikszentmihalyi, Applications of flow in human development and education (Springer, Dordrecht, 2014), 10.1007/978-94-017-9094-9.

5. F. Pels, J. Kleinert, F. Mennigen, Group flow: A scoping review of definitions, theoretical approaches, measures and findings. PLoS One 13, e0210117 (2018).

6. E. Hart, Z. Di Blasi, Combined flow in musical jam sessions: A pilot qualitative study. Psychology of Music 43, 275–290 (2013).

7. C. J. Walker, Experiencing flow: Is doing it together better than doing it alone? The Journal of Positive Psychology 5, 3–11 (2010).

8. R. K. Sawyer, Group genius : the creative power of collaboration (Basic Books, New York, ed. Revised edition., 2007), pp. xvi, 339 pages.

9. R. Hari, L. Henriksson, S. Malinen, L. Parkkonen, Centrality of Social Interaction in Human Brain Function. Neuron 88, 181–193 (2015).

10. M. Salanova, A. M. Rodriguez-Sanchez, W. B. Schaufeli, E. Cifre, Flowing together: a longitudinal study of collective efficacy and collective flow among workgroups. J Psychol 148, 435–455 (2014).

11. J. R. Katzenbach, D. K. Smith, The discipline of teams. Harv Bus Rev 71, 111–120 (1993).

12. J. R. Katzenbach, D. K. Smith, The wisdom of teams : creating the high-performance organization (Harvard Business School Press, Boston, Mass., 1993), pp. xii, 291 p.

13. I. Sato, “Bosozoku: flow in Japanese motorcycle gangs” in Optimal Experience: Psychological Studies of Flow in Consciousness, I. S. Csikszentmihalyi, M. Csikszentmihalyi, Eds. (Cambridge University Press, Cambridge, 1988), DOI: 10.1017/CBO9780511621956.006, pp. 92–117.

14. D. A. Stanley, R. Adolphs, Toward a neural basis for social behavior. Neuron 80, 816–826 (2013).

15. D. Y. Yang, G. Rosenblau, C. Keifer, K. A. Pelphrey, An integrative neural model of social perception, action observation, and theory of mind. Neurosci Biobehav Rev 51, 263–275 (2015).

16. D. Ongur, J. L. Price, The organization of networks within the orbital and medial prefrontal cortex of rats, monkeys and humans. Cereb Cortex 10, 206–219 (2000).

17. C. Lamm, J. Decety, T. Singer, Meta-analytic evidence for common and distinct neural networks associated with directly experienced pain and empathy for pain. Neuroimage 54, 2492–2502 (2011).

18. P. Molenberghs, R. Cunnington, J. B. Mattingley, Brain regions with mirror properties: a meta-analysis of 125 human fMRI studies. Neurosci Biobehav Rev 36, 341–349 (2012).

19. D. Dodell-Feder, J. Koster-Hale, M. Bedny, R. Saxe, fMRI item analysis in a theory of mind task. Neuroimage 55, 705–712 (2011).

20. M. Klasen, R. Weber, T. T. Kircher, K. A. Mathiak, K. Mathiak, Neural contributions to flow experience during video game playing. Soc Cogn Affect Neurosci 7, 485–495 (2012).

21. M. Ulrich, J. Keller, K. Hoenig, C. Waller, G. Gron, Neural correlates of experimentally induced flow experiences. Neuroimage 86, 194–202 (2014).

22. M. Ulrich, J. Keller, G. Gron, Dorsal Raphe Nucleus Down-Regulates Medial Prefrontal Cortex during Experience of Flow. Front Behav Neurosci 10, 169 (2016).

23. M. Ulrich, J. Keller, G. Gron, Neural signatures of experimentally induced flow experiences identified in a typical fMRI block design with BOLD imaging. Soc Cogn Affect Neurosci 11, 496–507 (2016).

24. D. J. Harris, S. J. Vine, M. R. Wilson, Neurocognitive mechanisms of the flow state. Prog Brain Res 234, 221–243 (2017).

25. T. W. Picton, S. A. Hillyard, Human auditory evoked potentials. II. Effects of attention. Electroencephalogr Clin Neurophysiol 36, 191–199 (1974).

26. H. E. Wang et al., A systematic framework for functional connectivity measures. Front Neurosci 8, 405 (2014).

27. G. Tononi, An information integration theory of consciousness. BMC Neurosci 5, 42 (2004).

28. M. Oizumi, L. Albantakis, G. Tononi, From the phenomenology to the mechanisms of consciousness: Integrated Information Theory 3.0. PLoS Comput Biol 10, e1003588 (2014).

29. P. J. Uhlhaas, W. Singer, Neural synchrony in brain disorders: relevance for cognitive dysfunctions and pathophysiology. Neuron 52, 155–168 (2006).

30. D. P. Kennedy, R. Adolphs, The social brain in psychiatric and neurological disorders. Trends Cogn Sci 16, 559–572 (2012).

31. H. H. Tseng et al., A systematic review of multisensory cognitive-affective integration in schizophrenia. Neurosci Biobehav Rev 55, 444–452 (2015).

32. G. Dumas, J. Nadel, R. Soussignan, J. Martinerie, L. Garnero, Inter-brain synchronization during social interaction. PLoS One 5, e12166 (2010).

33. J. Sanger, V. Muller, U. Lindenberger, Intra- and interbrain synchronization and network properties when playing guitar in duets. Front Hum Neurosci 6 (2012).

34. K. Yun, K. Watanabe, S. Shimojo, Interpersonal body and neural synchronization as a marker of implicit social interaction. Sci Rep 2, 959 (2012).

35. S. Dikker et al., Brain-to-Brain Synchrony Tracks Real-World Dynamic Group Interactions in the Classroom. Curr Biol 27, 1375–1380 (2017).

36. P. Goldstein, I. Weissman-Fogel, G. Dumas, S. G. Shamay-Tsoory, Brain-to-brain coupling during handholding is associated with pain reduction. Proc Natl Acad Sci U S A 115, E2528–E2537 (2018).

37. W. Zhang, M. M. Yartsev, Correlated Neural Activity across the Brains of Socially Interacting Bats. Cell 178, 413–428 e422 (2019).

38. K. L. Lu, H. Xue, T. Nozawa, N. Hao, Cooperation Makes a Group be More Creative. Cereb Cortex 29, 3457–3470 (2019).

39. U. Lindenberger, S. C. Li, W. Gruber, V. Muller, Brains swinging in concert: cortical phase synchronization while playing guitar. BMC Neurosci 10, 22 (2009).

40. M. Kawasaki, Y. Yamada, Y. Ushiku, E. Miyauchi, Y. Yamaguchi, Inter-brain synchronization during coordination of speech rhythm in human-to-human social interaction. Sci Rep 3, 1692 (2013).

41. H. Poikonen, P. Toiviainen, M. Tervaniemi, Naturalistic music and dance: Cortical phase synchrony in musicians and dancers. PLoS One 13, e0196065 (2018).

42. D. Engel, T. W. Malone, Integrated information as a metric for group interaction. Plos One 13 (2018).

43. E. Redcay, L. Schilbach, Using second-person neuroscience to elucidate the mechanisms of social interaction. Nat Rev Neurosci 20, 495–505 (2019).

44. C. Koch, G. Tononi, Can a Photodiode Be Conscious? New York Rev Books 60, 43–43 (2013).

45. H. Nolan, R. Whelan, R. Reilly, FASTER: fully automated statistical thresholding for EEG artifact rejection. Journal of neuroscience methods 192, 152–162 (2010).

46. C. Brunner, A. Delorme, S. Makeig, Eeglab - an Open Source Matlab Toolbox for Electrophysiological Research. Biomed Tech (Berl) 58 Suppl 1 (2013).

47. J. van Driel, T. Knapen, D. M. van Es, M. X. Cohen, Interregional alpha-band synchrony supports temporal cross-modal integration. Neuroimage 101, 404–415 (2014).

48. M. Stropahl, A. R. Bauer, S. Debener, M. G. Bleichner, Source-Modeling Auditory Processes of EEG Data Using EEGLAB and Brainstorm. Front Neurosci 12, 309 (2018).

49. P. E. Clayson, S. A. Baldwin, M. J. Larson, How does noise affect amplitude and latency measurement of event-related potentials (ERPs)? A methodological critique and simulation study. Psychophysiology 50, 174–186 (2013).

50. M. Reuter, N. J. Schmansky, H. D. Rosas, B. Fischl, Within-subject template estimation for unbiased longitudinal image analysis. Neuroimage 61, 1402–1418 (2012).

51. F. Tadel, S. Baillet, J. C. Mosher, D. Pantazis, R. M. Leahy, Brainstorm: a user-friendly application for MEG/EEG analysis. Comput Intell Neurosci 2011, 879716 (2011).

52. A. Gramfort, T. Papadopoulo, E. Olivi, M. Clerc, OpenMEEG: opensource software for quasistatic bioelectromagnetics. Biomed Eng Online 9, 45 (2010).

53. C. Destrieux, B. Fischl, A. Dale, E. Halgren, Automatic parcellation of human cortical gyri and sulci using standard anatomical nomenclature. Neuroimage 53, 1–15 (2010).

54. P. D. Welch, Use of Fast Fourier Transform for Estimation of Power Spectra - a Method Based on Time Averaging over Short Modified Periodograms. Ieee Transactions on Audio and Electroacoustics Au 15, 70–+ (1967).

55. M. Volker et al., The dynamics of error processing in the human brain as reflected by high-gamma activity in noninvasive and intracranial EEG. Neuroimage 173, 564–579 (2018).

56. A. Abraham et al., Machine learning for neuroimaging with scikit-learn. Front Neuroinform 8, 14 (2014).

57. A. Delorme et al., EEGLAB, SIFT, NFT, BCILAB, and ERICA: new tools for advanced EEG processing. Comput Intell Neurosci 2011, 130714 (2011).

58. L. A. Baccala, K. Sameshima, Partial directed coherence: a new concept in neural structure determination. Biol Cybern 84, 463–474 (2001).

59. A. Korzeniewska, M. Manczak, M. Kaminski, K. J. Blinowska, S. Kasicki, Determination of information flow direction among brain structures by a modified directed transfer function (dDTF) method. J Neurosci Methods 125, 195–207 (2003).

60. J. Geweke, Measurement of linear dependence and feedback between multiple time series. Journal of the American Statistical Association, 304–313 (1982).

61. S. L. Bressler, C. G. Richter, Y. Chen, M. Ding, Cortical functional network organization from autoregressive modeling of local field potential oscillations. Stat Med 26, 3875–3885 (2007).

62. W. G. P. Mayner et al., PyPhi: A toolbox for integrated information theory. PLoS Comput Biol 14, e1006343 (2018).

63. J. P. Lachaux, E. Rodriguez, J. Martinerie, F. J. Varela, Measuring phase synchrony in brain signals. Hum Brain Mapp 8, 194–208 (1999).

64. M. X. Cohen, Analyzing neural time series data : theory and practice, Issues in clinical and cognitive neuropsychology (The MIT Press, Cambridge, Massachusetts, 2014), pp. xviii, 578 pages, 516 unnumbered pages of plates.

