## Supplemental Information for "Team flow is a unique brain state associated with enhanced information integration and neural synchrony"

**This PDF file includes:**

- Supplementary text
- Figures S1 to S7
- Tables S1 to S3
- Legend for Movie S1

**Other supplementary materials for this manuscript include the following:**

- Movie S1

### Supplementary Method

#### Task

We used a commercial music rhythm game called "O2JAM U" (version 1.6.0.11, MOMO Co., South Korea) running on iPad air (model No. MD786LL/B, system, iOS 10.3.2). The basic structure of this game follows the most common structure in the music rhythm genre. Two consecutive screenshots of the game are shown in Fig. 1a. Visual cues (notes) moves in lanes from the top to the bottom of the screen where the tapping area is located. There are two kinds of cues: short and long ones. Players' task is to tap when a short cue reaches the tapping area, and to tap and hold for the duration a long cue is at the tapping area. The cues are designed to follow the rhythm and beats of music. This design gives the impression of playing a musical instrument thus produces much of the positive experience of the game. The game displays two types of real time feedback on the players' performance. The first feedback type includes a semantic judgement expression ("EXCELLENT", "GOOD", or "MISS") together with a numerical score presented at the center and the top corners of the screen (Fig. S1a). We made the first feedback type invisible to the participants, using a privacy screen protector, to enhance participants' focus on the tapping area. The second feedback type is a flashing visual effect that appears at the tapping area each time the player taps at the correct timing with the cue (Fig. S1a). We kept this feedback type visible to the participants as a reinforcement (Movie S1). The game provides two modes of play; a 2-lane or 4-lane modes where 2 or 4 lanes of moving cues are presented. We used the 2-lane mode during individual screening with the participant responsible for both lanes. We used the 4-lane mode during the main experiment where a pair of participants played with each participant responsible for two adjacent lanes.

After playing each song (trial), the game displays a performance report on the screen, including a final numerical score, the total number of cues, and the number of missed cues. The performance report of each trial was hidden from the participants until they finish answering their subjective experience questionnaire. The percentage of the missed cues per the total number of cues was used as a metric for the performance of each pair of participants.

#### Manipulations

To create the interpersonal synchronized audio-visual (Inter-SyncA) condition, the iPad was tilted and positioned, using a custom-made holder, equidistant from the pair of participants. Participants were instructed to sit on two chairs at a fixed distance, to keep their heads on chin rests, and to minimize their body movements except for fingers movement. The iPad was connected to a pair of stereo speakers placed horizontally and equidistant from the iPad and the pair of participants. A pair of participants played in the 4-lane mode.

Previous studies controlled the flow experience by manipulating the skill-challenge level in the experimental setup, either passively by free task performance and retrograde classification of certain time periods into several flow levels (20) or by actively controlling the task level to be either too easy (boredom), adaptive (flow), or too hard (overload) (21-23). One of the issues with modifying the skill-challenge level to study flow is that this changes other cognitive functions, such as attention, sensory information processing, and mental effort necessary to perform each task, as well as gross changes in motor behavior. Therefore, manipulating the skill-challenge level complicates distinguishing the neural mechanisms underlying team flow, individual flow, and team interaction. To avoid this complexity, we kept the task identical in all conditions using the same sequence of tapping cues. Instead, we manipulated the intrinsic reward/enjoyment dimension of flow by scrambling the game music and hence disrupting the synchronization between with the tapping cues and music rhythm (Table S1).

To create the interpersonal scrambled audio (Inter-ScrA) condition, participants played the same song (i.e. identical sequence of moving cues) as the Inter-SyncA using the same setup; however, a reversed and shuffled version of the music was played from the same speakers (Table S1). The music for each song was reversed through an online audio editing website (https://audiotrimmer.com/online-mp3-reverser/) and then cut into 5-second fragments through an online audio cutter (http://mp3cut.net/). We randomly shuffled the fragments and rejoined them through an online audio joiner (http://audio-joiner.com/).

To create the occlusion synchronized audio-visual (Occl-SyncA) condition, participants played the same song (i.e. identical sequence of moving cues) as the Inter-SyncA using the same setup; however, a black foam board (1 cm x 1.5 m x 75 cm) was placed between the chairs to completely block the participants' view of each other and a black card board was placed across the iPad screen with an opening to show the visual cues but not the tapping area (Fig. S1a and Table S1).

**Screening**

First, participants were tested using a selected song (269 cues per the 2 lanes) in the 2-lane mode to exclude participants novice to the game. Participants were qualified to complete the study if they missed no more than 10 cues. Out of the 78 participants recruited for the first screening test 54 participants were qualified. Qualified participants were also tested using other selected songs with higher number of cues to check their skill level. Because the positive experience of this game depends on individual preference for the music rhythm, we prepared 11 songs (500 to 960 cues per the 4 lanes) in various genres. Qualified participants were asked to rate their preferences for all the 11 songs separately on a 7-point scale (1 for "not like it at all", 7 for "like very much"). Although the duration of the songs varied from 110 to 160 seconds, each song was played only for 1 minute, which was long enough to ensure that participants had heard the main rhythm of the song. We fixed duration for all songs to ensure the accuracy of preference rating. To avoid possible influence from experimenter, the experimenter held a neutral attitude with no body movements syncing with the music (e.g., nodding, taping fingers or feet, etc.) or eye contact with participants. Participants were paired based on their skill level and song preference rating.

Second, we screened paired participants for their preference to the interpersonal set-up (Inter-SyncA) versus playing in the occlusion set-up (Occl-SyncA ) or a solo set-up (Solo-SyncA) in a behavior pilot experiment. For the Solo-SyncA condition, we used the 2-lane mode of the game. At the end of the pilot experiment, participants rated their preference on each condition on a scale from 1 to 7. Participants were excluded if they rated either of the Occl-SyncA or Solo-SyncA higher than the Inter-SyncA condition. Unexcluded pro-social participants (38 participants) were invited for the main experiment on a separate day. For both the screening and the main experiment, we paired only same gender participants and the main pairing criteria for qualified participants was their skill level and song preferences. As friends were encouraged to pair up to participate in this experiment, we preferred pairing friends into the main experiment if they satisfy the main criteria.

**Main experiment**

After setting the EEG cap, the electrode positions were co-registered with the T1-MRI images using Brainsight TMS Navigation system (Rogue Resolutions Ltd, United Kingdom). Then paired participants, seated on two chairs at a fixed distance, underwent a beep-only trial. In this trial, they were instructed to passively listen to the task-irrelevant beep stimulus for 2.5 minutes, to keep their heads on chin rests and their eyes open, and to minimize their body movements. This trial was to check the EEG recording quality and verify that we could obtain a clear AEP response. Then paired participants performed a practice trial in the Occl-SyncA condition to get familiarized with the procedure. Then each pair of participants was required to play 6 songs each at the Inter-SyncA, Inter-ScrA, or Occl-SyncA conditions forming 18 trials (Fig. S1b,c). One pair played only 5 songs due to time availability. The sequence of songs and conditions were pseudo-randomized (Fig. S1c). To keep participants' continuous interest, the consecutive songs were always different (Fig. S1c). To control practice and carryover effects, we arranged each condition to have equal chance of being before or after the other 2 conditions (Fig. S1c). All trials included a resting phase and a playing phase (Fig. S1b). During the resting phase, participants were instructed to passively listen to the task-irrelevant beep sound and the game background music for 30 seconds, to keep their heads on chin rests and their eyes open, and to minimize their body movements. Then, the experimenter asked participants to click the game-play icon on the iPad to start the playing phase.

During the playing phase, participants were instructed to keep their heads on chin rests, to minimize their body movements except for finger movement, and to minimize vocal sounds that could distract their partner. They were allowed to give verbal comments related to the game. We video recorded a top-view of the iPad and the participants' hands using an iPhone fixed approximately 50 cm above the iPad where all the types of feedbacks were visible. After the playing phase of each trial, participants were given access to private screen and keyboards to freely answer the questionnaires on the flow experience and team interaction experience (Fig. S1c and Fig. S2). Then, the pair were allowed to jointly view the performance report. The experimenter asked the participants whether they wanted to proceed to the next trial or if they needed some rest. This is to minimize the effect of fatigue on performance or EEG recording quality.

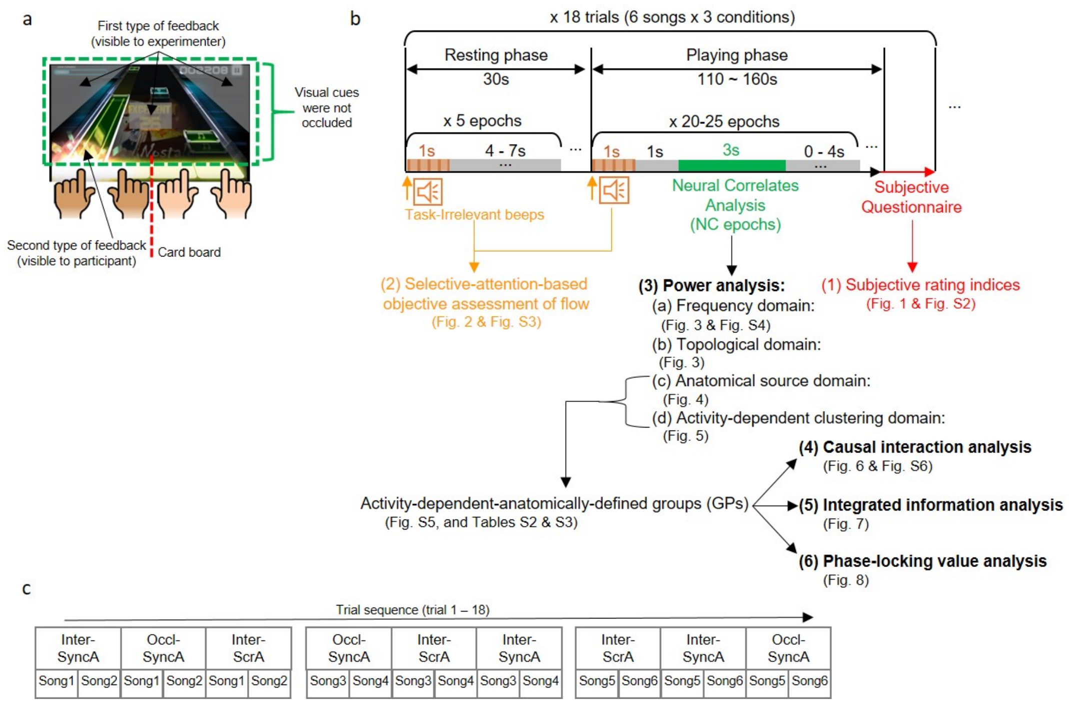


**Figure S1. Experimental design**

**a**, A screen shot of the game showing feedbacks and position of the card board. The first type of feedback was invisible to participants. The second type of feedback was visible to participants in the Inter-SyncA and Inter-ScrA conditions. The card board (red dashed line) occluded the tapping area, the second type of feedback, the whole bodies of participants including fingers. The card board kept the visual cues visible to participants (green dashed rectangle). **b**, Trial and analyses details: Participants were sitting still while listening to a background music during the resting phase. The electroencephalogram was epoched for the auditory-evoked potential (AEP) analysis of the task-irrelevant beep sound (AEP epochs; orange) and for the neural correlates analysis (NC epochs; green). All neural correlate analyses, with the corresponding figure or table, are summarized in steps (3)-(6). **c**, Sequence of the 18 trials, showing which song and condition per trial was assigned, during the main experiment.

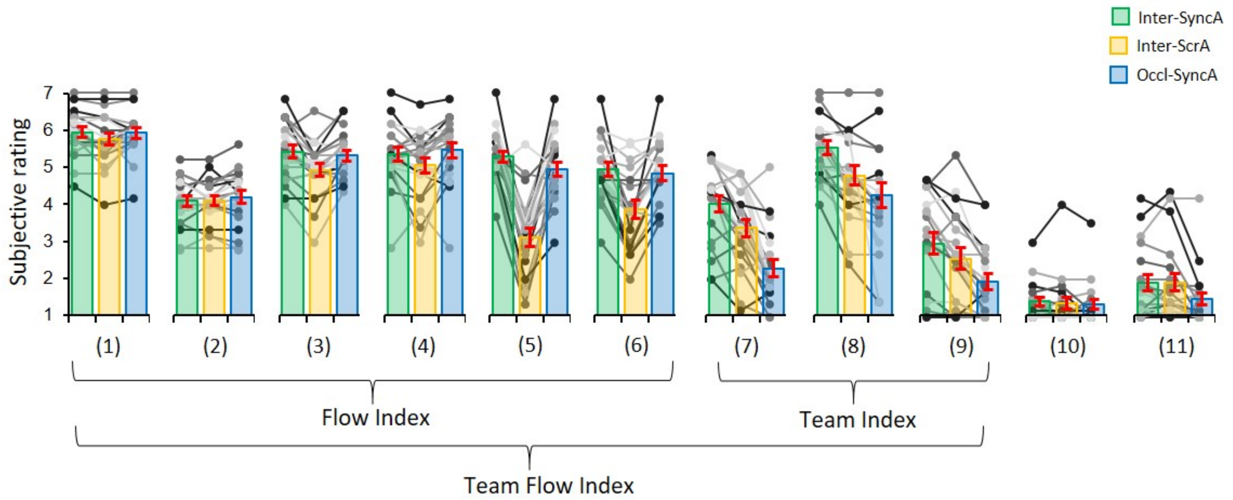


**Figure S2. Summary of subjective ratings for assessing the flow state and team interaction.** The flow index was calculated by averaging ratings for questions 1 to 6, the team index was calculated by averaging ratings for questions 7 to 9, and the team flow index was calculated by averaging ratings for questions 1 to 9. Question description: (1) “I had the necessary skill to play this trial successfully.”; (2) “I will enjoy this trial more if it has less/more notes.”; (3) “I felt in control while playing this trial.”; (4) “I made correct movements automatically without thinking.”; (5) “I love the feeling of this trial and want to play it again.”; (6) “How time flies during this trial?”; (7) “I was aware of the other player's actions.”; (8) “I felt like I was playing with the other person as a team.”; (9) “I was coordinating my fingers with the other player’s fingers.”; (10) “I felt like I was competing with the other player.”; (11) “I was distracted by the other player’s actions.”; [for question (2), rating 7 = more notes and 1 = less notes; for question (6), rating 7 = fast and 1 = slow; for other questions, rating 7 = strongly agree and 1 = strongly disagree]. Error bars represent mean ± s.e.m.; n = 20.

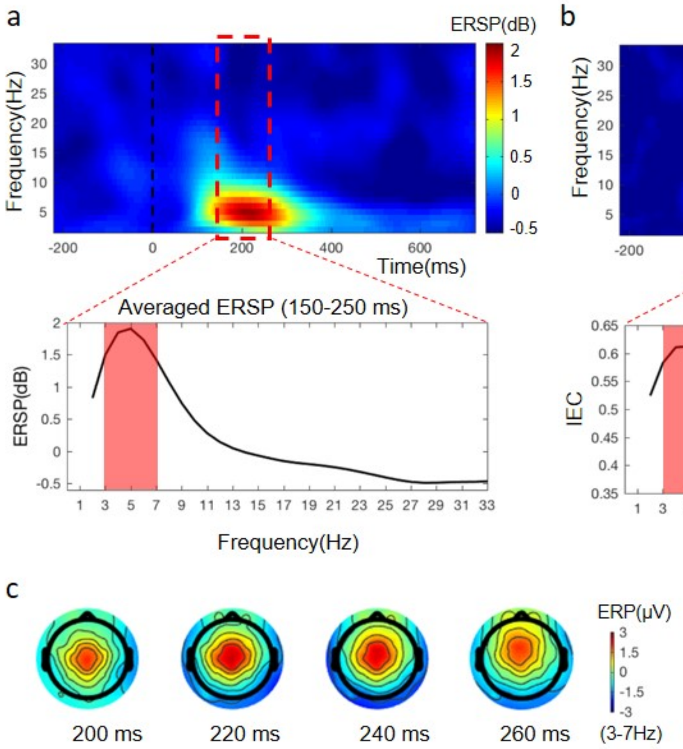


**Figure S3.** **a**,**b**, Time-frequency analysis of the auditory-evoked potential (AEP) locked to the task-irrelevant beep onsets presented as the event-related spectral perturbation (ERSP) in A and the inter-epoch coherence analysis (IEC) in B. Both ERSP and ITC showed changes in theta activity at 100 – 350 ms post-onset (upper panel). An increase in theta activity (3 – 7 Hz) was prominent at 150 – 250 ms post-onset (lower panel). **c**, Topographies of the event-related potential (ERP), bandpass-filtered in the theta range (3 – 7 Hz), at the indicated timepoints (ms) from the task-irrelevant beeps showing enhanced potential at the central channels.

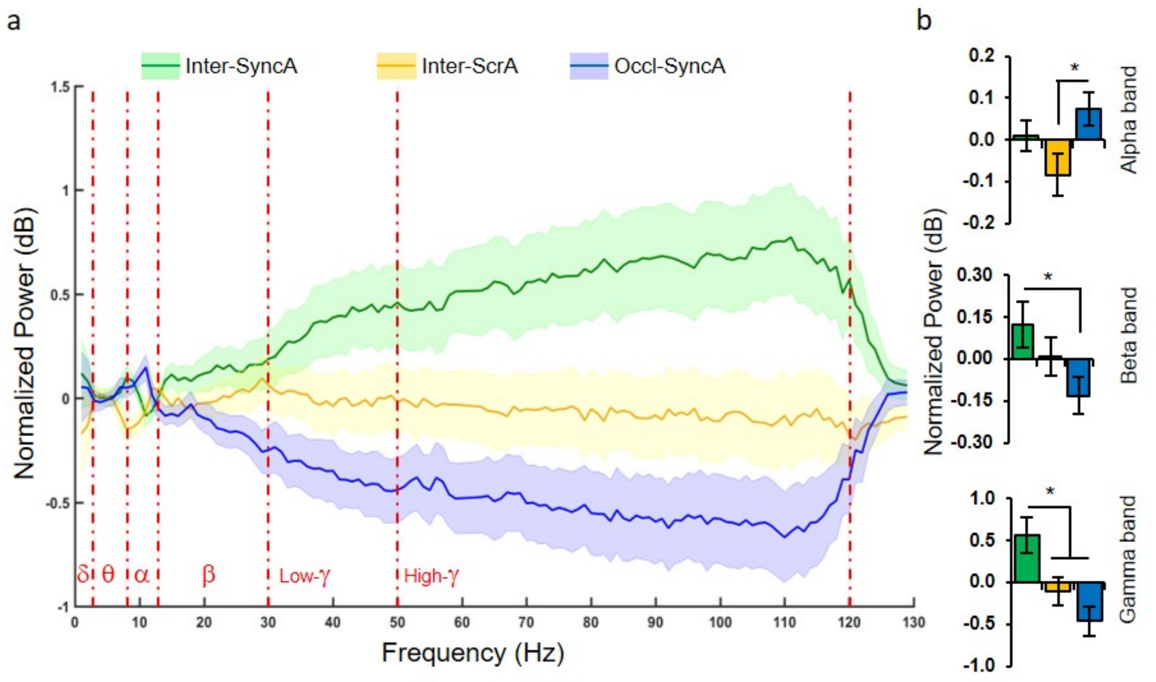


**Figure S4. a**, The power difference spectral analysis for the three conditions grand averaged for all the 128 channels. **b**, Averaged individual power difference for the alpha (8 – 12 Hz), the beta (13 – 30 Hz) and gamma (31 – 120 Hz) frequency bands. One-way ANOVA with Bonferroni post-hoc test. * $P < 0.05$. Error bars represent mean ± s.e.m.; $n = 20$.

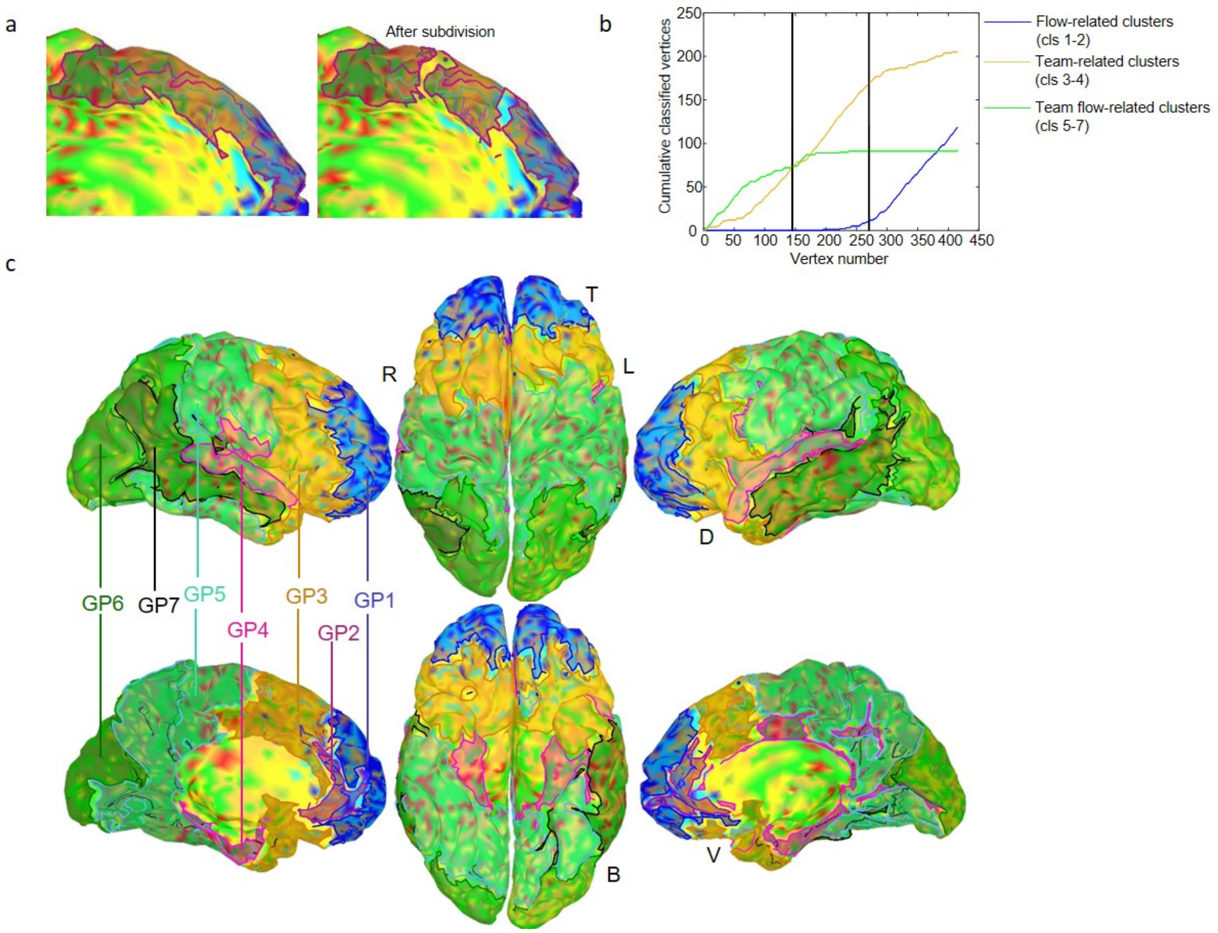


**Figure S5. Activity-dependent anatomically-defined grouping of ROIs (GPs). a,** A medial view of the left superior frontal cortex (area inside the red boundary/transparent contour) before (left panel) and after (right panel) subdivision. **b**, The cumulative cluster composition curve for the left superior frontal cortex. The subdivision thresholds are shown as two vertical black lines subdividing this ROI into three subdivisions: flow-related subdivision (cls1-2), team-related subdivision (cls3-4), and team flow-related subdivision (cls5-7). **c**, Transparent contours showing the activity-dependent-anatomically-defined groups (GP1-7) which is also summarized in Table S3. B, bottom; D, dorsal; L, left; R, right; T, top; V, ventral.

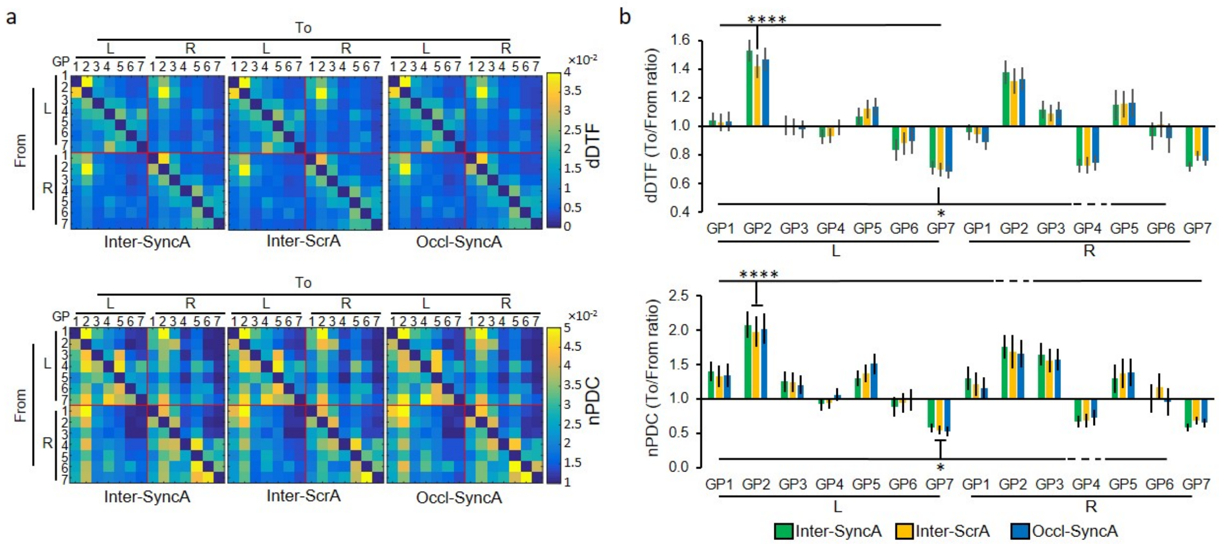


**Figure S6. Information flow analysis showing the middle temporal group (GP7) receives information from other groups. a**, The mean direct directed transfer function (dDTF) and normalized partial direct coherence (nPDC) causal interaction matrices for the activity-dependent-anatomically-defined groups (GPs). To, indicates sending information; From, indicates receiving information; L, left hemisphere; R, right hemisphere. **b**, The mean To/From ratio for dDTF and nPDC. In both measures, The To/From ratio showed L-GP2 as a significant global information sender and L-GP7 as a significant global information receiver. Two-way ANOVA with Tukey's post hoc test. * $P < 0.05$, **** $P < 0.0001$. Dashed line indicates $p > 0.05$. Error bars represent mean ± s.e.m.; n = 20.

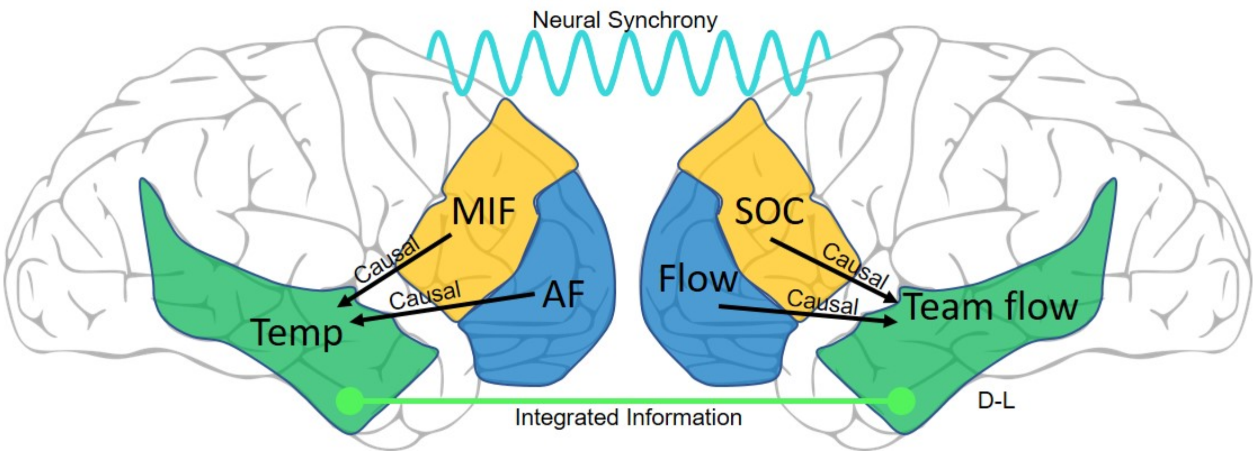


**Figure S7. A neural model for team flow.** The temporal cortex (Temp, green region) is uniquely activated during team flow (check Fig. 3, 4, 5, and Fig. S5). The anterior frontal cortex (AF, blue region), and the middle and inferior frontal cortex (MIF, yellow region) are uniquely activated during individual flow (Flow) and social interaction (SOC), respectively (check Fig. 5, and Fig. S5). During team flow, Left Temp causally receives more information (black arrows) from the contralateral, shown here as ipsilateral for simplicity, AF and MIF (check Fig. 6, and Fig. S6). Left Temp is significantly involved in higher inter-brain integrated information (green line) during team flow (check Fig. 7). Higher inter-brain neural synchrony (tan wavy line) during team flow (check Fig. 8).

**Table S1.**
**Behavioral manipulation comparison.**

| *Condition names* | | **Inter-SyncA** | **Inter**-ScrA | Occl-**SyncA** |
|---|---|---|---|---|
| *Predicated outcome* | | **Team**<br>**flow** | **Team**<br>non-flow | Non-Team<br>**flow** |
| *Cues sequence (visual stimulus)* | | Fixed (across time) | | |
| *Song (auditory stimulus)* | | Fixed (across trials) | | |
| | | Synced with cues | Scrambled | Synced with cues |
| *Task-irrelevant stimulus sequence* | | Fixed (pseudorandomized) | | |
| *Partner's* | *Cues* | Visible/fixed (across time) | | |
| | *Feedback (Flashes)* | Visible/Dynamic | Visible/Dynamic | Blocked |
| | *Finger/body* | Visible/Dynamic | Visible/Dynamic | Blocked |

**Table S2.**
**Cluster composition of the activity-dependent anatomically-defined groups (GPs).**

| | cls1 | cls2 | cls6 | cls7 | cls3 | cls4 | cls5 | Flow -related | Team -related | Team flow-related |
|---|---|---|---|---|---|---|---|---|---|---|
| **L-GP1** | 36.84 | 27.76 | 16.53 | 18.28 | 0.00 | 0.00 | 0.60 | **64.61** | 34.80 | 0.60 |
| **R-GP1** | 35.99 | 31.30 | 16.65 | 15.79 | 0.00 | 0.00 | 0.27 | **67.29** | 32.44 | 0.27 |
| **L-GP2** | 31.13 | 16.56 | 16.56 | 35.10 | 0.66 | 0.00 | 0.00 | **47.69** | 0.66 | **51.66** |
| **R-GP2** | 26.11 | 20.38 | 28.66 | 24.84 | 0.00 | 0.00 | 0.00 | **46.49** | **53.50** | 0.00 |
| **L-GP3** | 8.58 | 1.69 | 23.50 | 51.68 | 10.59 | 0.10 | 3.86 | 10.27 | **75.18** | 14.55 |
| **R-GP3** | 8.94 | 2.63 | 40.49 | 36.82 | 8.82 | 0.99 | 1.30 | 11.58 | **77.31** | 11.11 |
| **L-GP4** | 0.36 | 0.12 | 25.85 | 23.63 | 38.11 | 6.17 | 5.75 | 0.49 | **49.48** | **50.04** |
| **R-GP4** | 0.00 | 0.00 | 31.48 | 21.04 | 36.67 | 8.65 | 2.15 | 0.00 | **52.52** | **47.48** |
| **L-GP5** | 0.00 | 0.00 | 16.48 | 12.24 | 53.02 | 8.27 | 10.00 | 0.00 | 28.72 | **71.29** |
| **R-GP5** | 0.00 | 0.11 | 16.14 | 11.14 | 51.85 | 8.93 | 11.84 | 0.11 | 27.27 | **72.62** |
| **L-GP6** | 0.00 | 0.00 | 20.58 | 24.40 | 38.52 | 0.81 | 15.68 | 0.00 | **44.99** | **55.02** |
| **R-GP6** | 0.05 | 0.00 | 0.73 | 8.79 | 35.15 | 0.64 | 54.63 | 0.05 | 9.52 | **90.43** |
| **L-GP7** | 0.00 | 0.00 | 18.08 | 11.91 | 49.45 | 7.68 | 12.87 | 0.00 | 30.00 | **70.00** |
| **R-GP7** | 0.00 | 0.00 | 12.13 | 10.33 | 46.84 | 8.35 | 22.35 | 0.00 | 22.46 | **77.54** |

**Table S3.**
**Anatomical composition of the activity-dependent anatomically-defined groups (GPs).**

| | Anatomical ROIs (left hemisphere) | Anatomical ROIs (right hemisphere) |
|---|---|---|
| GP1 | S_front_middle L; S_suborbital L; G&S_frontomargin L; G&S_transv_frontopol L | G&S_frontomargin R; S_front_middle R; G&S_transv_frontopol R |
| GP2 | G&S_cingul-Ant L | G&S_cingul-Ant R |
| GP3 | S_temporal_transverse L; S_orbital_lateral L; Pole_temporal L; Lat_Fis-ant-Vertical L; G&S_cingul-Mid-Ant L; S_orbital-H_Shaped L; S_circular_insula_ant L; G_front_inf-Opercular L; S_orbital_med-olfact L; G_temp_sup-Plan_polar L; S_front_inf L; G_subcallosal L; G_front_inf-Orbital L; G_front_inf-Triangul L; Lat_Fis-ant-Horizont L | G&S_cingul-Mid-Ant R; G&S_cingul-Mid-Post R; S_precentral-inf-part R; G_subcallosal R; S_suborbital R; G_temp_sup-G_T_transv R; G_insular_short R; S_circular_insula_ant R; S_circular_insula_sup R; S_orbital-H_Shaped R; G_temp_sup-Plan_polar R; G_front_inf-Opercular R; G_front_inf-Orbital R; S_front_inf R; Pole_temporal R; S_orbital_med-olfact R; S_orbital_lateral R; G_front_inf-Triangul R; Lat_Fis-ant-Horizont R; Lat_Fis-ant-Vertical R |
| GP4 | S_circular_insula_sup L; S_precentral-inf-part L; S_subparietal L; G_oc-temp_med-Parahip L; G_insular_short L; G_temp_sup-Lateral L; S_collat_transv_ant L; G&S_cingul-Mid-Post L; G_Ins_lg&S_cent_ins L; S_pericallosal L; S_calcarine L; S_circular_insula_inf L | G_temp_sup-Lateral R; G&S_subcentral R; S_temporal_transverse R; G_Ins_lg&S_cent_ins R; G_oc-temp_med-Parahip R |
| GP5 | G_oc-temp_med-Lingual L; S_precentral-sup-part L; G&S_subcentral L; G_precuneus L; S_cingul-Marginalis L; G_oc-temp_lat-fusifor L; G_cingul-Post-dorsal L; G_temp_sup-G_T_transv L; G_precentral L; G&S_paracentral L; G_cingul-Post-ventral L; S_central L; G_pariet_inf-Supramar L; G_postcentral L; S_postcentral L; S_interm_prim-Jensen L | S_collat_transv_ant R; G_precentral R; S_central R; S_circular_insula_inf R; S_precentral-sup-part R; G_cingul-Post-ventral R; G_temporal_inf R; G_cingul-Post-dorsal R; S_oc-temp_med&Lingual R; G_postcentral R; S_oc-temp_lat R; G_oc-temp_lat-fusifor R; S_cingul-Marginalis R; G_oc-temp_med-Lingual R; G&S_paracentral R; S_postcentral R; S_subparietal R; G_precuneus R; G_pariet_inf-Supramar R; S_calcarine R; S_interm_prim-Jensen R |
| GP6 | Pole_occipital L; S_oc_middle&Lunatus L; G_occipital_middle L; G_cuneus L; G&S_occipital_inf L; S_parieto_occipital L; G_occipital_sup L; S_collat_transv_post L; S_oc_sup&transversal L; G_pariet_inf-Angular L; G_parietal_sup L; S_intrapariet&P_trans L | S_oc_sup&transversal R; G_occipital_middle R; G_occipital_sup R; Pole_occipital R; G_cuneus R; S_oc_middle&Lunatus R; G&S_occipital_inf R; G_parietal_sup R; S_collat_transv_post R; S_parieto_occipital R; S_intrapariet&P_trans R |
| GP7 | G_temp_sup-Plan_tempo L; S_occipital_ant L; S_oc-temp_lat L; S_oc-temp_med&Lingual L; Lat_Fis-post L; | G_pariet_inf-Angular R; S_occipital_ant R; S_temporal_sup R; G_temp_sup-Plan_tempo R; |

| | | |
|---|---|---|
| | S_temporal_sup L; S_temporal_inf L; G_temporal_inf L; G_temporal_middle L | S_temporal_inf R; Lat_Fis-post R; G_temporal_middle R |
| Subdivided | G_orbital L; G_rectus L; G_front_middle L; S_front_sup L; G_front_sup L | G_rectus R; G_orbital R; G_front_middle R; S_front_sup R; G_front_sup R; S_pericallosal R |

**Legend for Movie S1**

A few seconds of game-play in the Inter-SyncA and Inter-ScrA conditions. The "Beeps" word at the bottom right indicates the timing of the task-irrelevant beep sound presentation. These words are overlaid in the video for illustration and were not present during the experiment. The scores and other indicators at the center, and at the top right and left corners were hidden to the participants.
